# Herpes Simplex Virus Assembly and Spread in murine skin after infection from the outside

**DOI:** 10.1101/2024.09.18.613798

**Authors:** Timmy Richardo, Xiaokun Liu, Katinka Döhner, Anna Buch, Anne Binz, Anja Pohlmann, Madeleine de le Roi, Wolfgang Baumgärtner, Korbinian Brand, Rudolf Bauerfeind, Reinhold Förster, Beate Sodeik, Stephan Halle

## Abstract

Herpes simplex viruses (HSV) cause many skin diseases, particularly in immunocompromised patients. HSV-1 infection of murine skin recapitulates many aspects of human pathology. However, many protocols rely on mechanical or enzymatic skin disruption to induce lesions, although this can alter skin homeostasis and prime antiviral inflammation before inoculation. To investigate the initial events following HSV-1 primary skin infection before the onset of symptoms, we developed a novel murine *ex vivo* explant model using gentle depilation but no further scarification and infected keratinocytes from the outside with minimal tissue damage. Two-photon microscopy studies showed that HSV-1 spread exclusively in the epidermis. The infection centers increased in number and size over time and contained hundreds of infected keratinocytes. We investigated the HSV-1 spread at the cellular level, using reporter strains with fluorescently-tagged capsid protein VP26, and monitored the formation of nuclear capsid assembly sites, nuclear capsid egress, and the recruitment of the inner tegument protein pUL37GFP, the outer tegument protein VP11/12GFP, and the envelope protein gDGFP to cytoplasmic capsids. By electron microscopy, the skin appeared intact, and keratinocytes contained many nuclear capsids, primary virions in the nuclear envelope, cytosolic membrane-associated capsids, and enveloped virions. Our protocol provides a robust and reproducible approach to investigate the very early events of HSV-1 spread in the skin, to characterize the phenotypes of HSV-1 mutants in terminally differentiated skin tissues, and to evaluate potentially antiviral small molecules in a preclinical *ex vivo* infection model.

**IMPORTANCE:** This study describes a novel murine *ex vivo* skin explant model to investigate early events in HSV-1 infection without causing significant tissue damage. To infect from the outside, via the apical keratinocytes, this method relies on gentle depilation, which maintains skin integrity. HSV-1 spread exclusively within the epidermis, with infection centers growing over time and involving hundreds of keratinocytes. Using advanced microscopy techniques, we tracked HSV-1 spread at the cellular level and intracellular assembly of all intermediate virus structures. This model offers a valuable tool for studying the initial stages of HSV-1 infection, assessing viral mutant phenotypes, and testing antiviral compounds in a more physiological context to provide critical insights into HSV-1 pathogenesis and therapeutic strategies.

## INTRODUCTION

The WHO estimates that about two-thirds of the human population is infected with herpes simplex virus (HSV) type 1 and about 13% with HSV-2 (James et al., 2020; Looker et al., 2015). HSV-1 typically causes cold sores and lesions on the oral mucosa and the perioral skin, while HSV-2 can lead to genital skin lesions and ulcers (Singh et al., 2019; Whitley and Roizman, 2016). After infection from the outside and local replication, HSV-1 and HSV-2 enter neurites innervating the primary site of inoculation and establish lifelong latent infections in the neurons of the cranial and dorsal root ganglia (Cohen, 2020; Koyuncu et al., 2021). Upon reactivation in these latently infected neurons, progeny HSV particles return via anterograde transport and re-infect the skin or the mucosa from the inside.

Primary HSV-1 infections, as well as reactivations from latency, can lead to fatal encephalitis, blinding keratitis, disseminated disease, or eczema herpeticum (Traidl et al., 2021; Whitley and Roizman, 2016; Piret and Boivin, 2020; Tan et al., 2022; Wurzer et al., 2017). About 22% of patients with moderate to severe atopic dermatitis have a history of eczema herpeticum (Traidl et al., 2023, 2021; Weidinger et al., 2018). Moreover, the stigma associated with facial or genital lesions leads to psychological distress. Despite the availability of life-saving antiviral drugs like acyclovir, current treatments are still inadequate, in particular for immunocompromised patients as well as for the very young and the elderly; up to 70% of the patients surviving herpes simplex encephalitis maintain neurological symptoms (Gnann and Whitley, 2017; Piret and Boivin, 2020).

Healthy skin provides an effective barrier against pathogens (Coates et al., 2018; Nestle et al., 2009; Pasparakis et al., 2014). The epidermis is a tight epithelium composed of the layers stratum corneum, stratum granulosum, stratum spinosum, and stratum basale. It consists mainly of keratinocytes, which also line up the hair follicles, but also entails melanocytes, macrophages, Langerhans cells, and T cells. The thicker dermis underneath the epidermis consists mainly of fibroblasts and the extracellular collagen matrix and contains blood cells and lymphatic vessels. Nevertheless, HSV can infect the skin, and its replication in keratinocytes and fibroblasts leads to the development of herpetic lesions (Cunningham et al., 1985; Gebhardt et al., 2011; Hor et al., 2015; Kim et al., 2012).

Upon *ex vivo* infection of human genital skin biopsies, HSV-1 replication is limited predominantly to epidermal keratinocytes and mucosal epithelial cells but with little access to dermal fibroblasts, while *ex vivo* infection of isolated fibroblasts yields high titers of viral progeny (Kim et al., 2015). With the experimental protocols reported so far, HSV-1 infects abdominal skin explants from healthy donors only from the basolateral side after removal of the dermis and the basement membrane, but *ex vivo* infection of human oral epithelial cells is also productive (Bertram et al., 2021; Thier et al., 2017). Even mechanical wounding of adult human skin with microneedles is not sufficient to disrupt the protective skin barrier against HSV-1 infection from the outside (De La Cruz et al., 2021; Tajpara et al., 2019).

Although the anatomy and immunology of human and murine skin differ (Pasparakis et al., 2014), infection of murine skin recapitulates many aspects of human HSV-1 skin diseases (Hill and Shimeld, 1998; Kollias et al., 2015; Hussain et al., 2024; Tsalenchuck et al., 2014). Inoculation of mice after flank scarification results in local HSV-1 amplification and the formation of primary lesions that heal within approximately one week (Silva et al., 2017; van Lint et al., 2004, 2005). In such experiments, the skin homeostasis might be altered, and inflammatory responses activated before the infection as the epidermis and dermis are often dissociated to some extent by proteases, abrasion, or hair removal by waxing or tape-stripping (Amberg et al., 2017; Oyoshi et al., 2010; St Laurent et al., 2021; van Lint et al., 2004; Wen et al., 2022). These models have nevertheless provided important insights into the immune responses in the skin (Allan et al., 2003; Bedoui et al., 2009; Gebhardt et al., 2011; Wakim et al., 2008). Moreover, HSV-1 spread has been studied *ex vivo* in murine skin explants, but most studies have been focused on infecting the dermis, which recapitulates skin infection from the inside after reactivation from latency (Petermann et al., 2015; Puttur et al., 2010; Rahn et al., 2017, 2015; Stock et al., 2004).

Since the very early events of skin lesion formation before the onset of symptoms are not well understood, we have established a novel 3-dimensional experimental system to investigate the initiation of HSV-1 infection from the outside in terminally differentiated murine skin explants. Using confocal and two-photon fluorescence microscopy, we show that HSV-1 had formed infection centers already at 12 h post inoculation, and then spread laterally in terminally differentiated cells. Our HSV-1 dual-color reporter strains HSV1-CheVP26-pUL37GFP and HSV1-CheVP26-VP11/12GFP faithfully reflected the sequential acquisition of inner and outer tegument proteins onto capsids (reviewed in Döhner et al., 2024). Using those reporter strains with fluorescent tags on the capsid, the tegument, and the viral envelope, as well as electron microscopy, we show that all HSV-1 assembly intermediates were formed, that capsids underwent primary envelopment, recruited the inner tegument protein pUL37 and the outer tegument protein VP11/12 in the cytoplasm, and underwent secondary envelopment on cytoplasmic membranes containing the envelope glycoprotein gD. Our protocol using skin explant cultures bridges studies infecting primary cells cultured in dishes or on glass with animal infection experiments. It provides a versatile, cost-effective system to investigate cutaneous HSV-1 infection to characterize the phenotype of HSV-1 mutants and to test the potency of novel small-molecule candidates for antiviral therapy that likely could be expanded to other cutaneous viral and bacterial infections.

## RESULTS

### A novel *ex vivo* model for HSV-1 skin infection

To investigate the very early events of HSV-1 skin infection, we used explant cultures from the ears of six to nine weeks old mice. The hair was removed by a 5 min treatment with a depilatory cream (Fig. 1). The ears were then split into dorsal and ventral sheets and glued onto slightly larger tissue gaze pieces, which in turn were placed onto cell culture medium to generate an air-liquid-interface culture. Immunohistochemistry analyses of cross-sections showed that the depilation had removed not basal keratinocytes expressing keratin-14 expressing keratinocytes (Fig. 2, Aii, Bii; red in Aiv, Biv), but some apical keratinocytes as indicated by the reduction of loricrin (Fig. 2 Aiii, Biii; green in Aiv, Biv). Together with keratin 1 and keratin 10, loricrin is a major protein of the cornified keratinocytes (Menon et al., 2012; Wallace et al., 2012).

**Figure 1.**
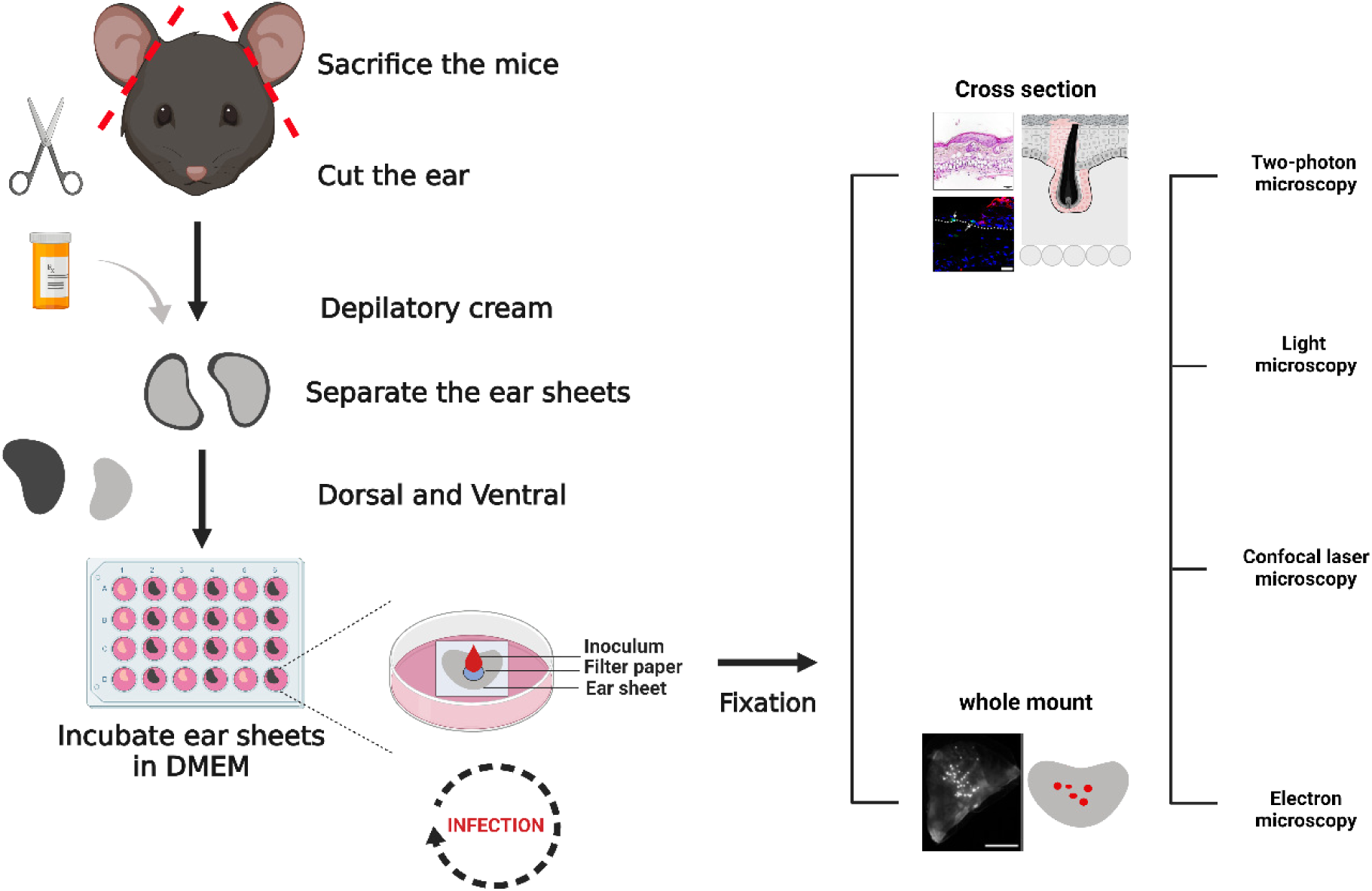
*Ex-vivo* HSV-1 infection of murine skin. The ears of sacrificed mice were collected and depilated. The entire dorsal and ventral ear sheets of about 1 cm^2^ were placed on a nylon gaze in an air-liquid-interface culture with the dermis side facing the medium. Filter papers with a diameter of 5 mm were positioned onto the epidermis surface, and 10 µL of buffer without or with HSV-1 were pipetted directly onto a given filter paper. After an incubation of 30 min at 37°C, the filter papers were removed, and the explants were cultured at 37°C for up to 96 h, fixed or lysed, and processed for microscopy, molecular biology, or virology assays. Semi-thin cryosections and paraffin-sections were labeled with primary antibodies against host or HSV-1 proteins followed by secondary fluorescent antibodies. HSV-1 infected cells were localized also by the expression of fluorescent proteins of the HSV-1 reporter strains. The specimens were analyzed by confocal or 2-photon fluorescence microscopy as well as by electron microscopy.

**Figure 2.**
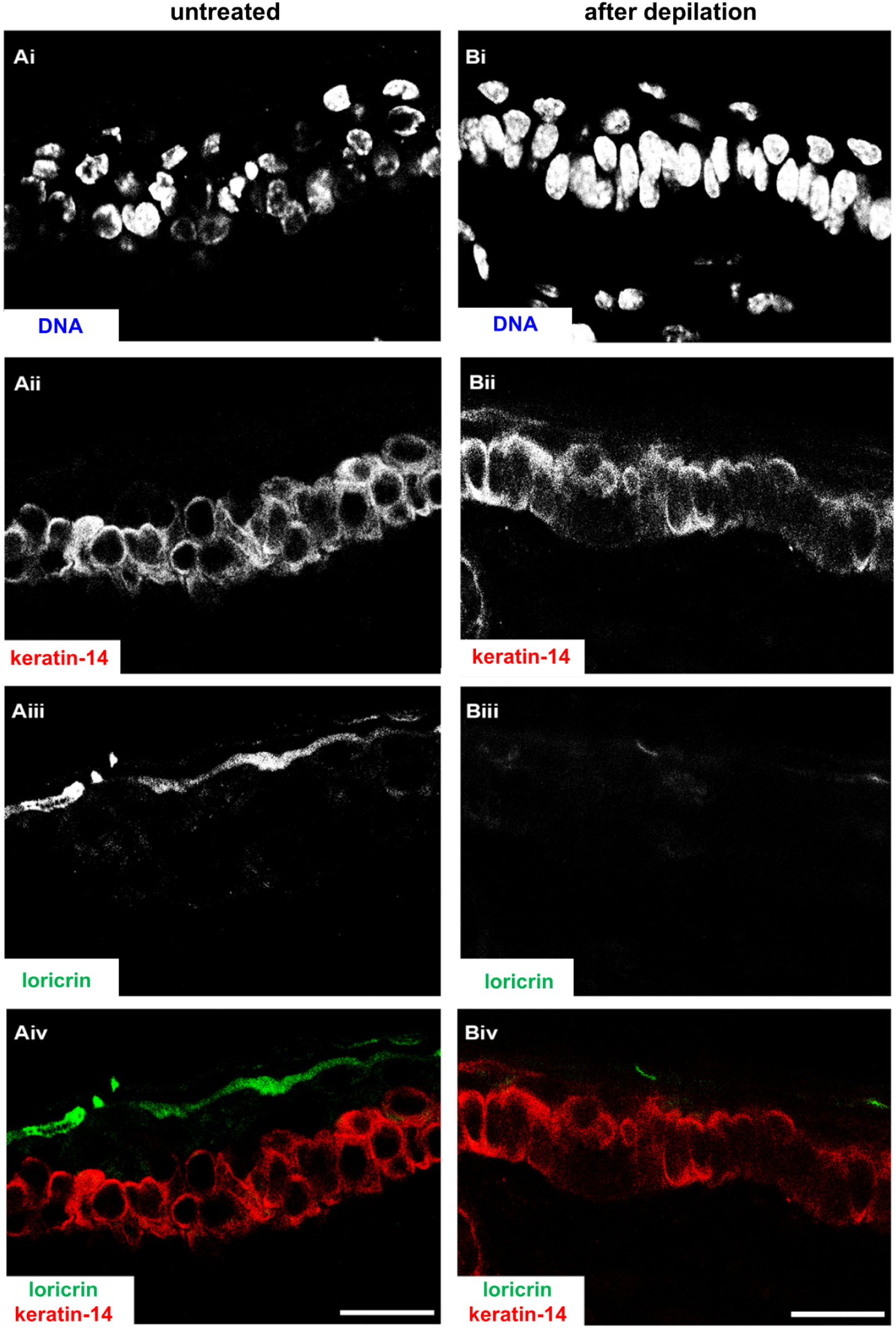
Depilation-induced changes in skin morphology. Uninfected murine ear sheets without (A), or after depilation (B) were cultured at the air-liquid interface for 48 h, fixed, and embedded. Cryosections were stained for DNA (DAPI; Ai, Bi), labeled for keratin-14 (Aii, Bii red in Aiv, Biv), and loricrin (Aiii,Biii green in Aiv, Biv), and analyzed by confocal fluorescence microscopy. Scale bar 25 µm.

For virus inoculation, round filter papers were placed onto the center of a given dorsal or ventral skin sheet, and a medium containing HSV1(17^+^)Lox-Che was added directly onto the filter papers (Fig. 1; Table 1). The reporter strain HSV1-Che expresses the fluorescent reporter mCherry under the control of a constitutively active viral promoter (Krawczyk et al., 2015). After 30 min, the filter papers were removed, and the explants were transferred back to a 37°C incubator for up to 96 h. Within 48 h post-infection (hpi), HSV-1 genomes (Fig. 3A) and HSV-1 transcripts (Fig. 3B) were clearly detected in infected samples. The HSV-1 gene UL27, encoding for the glycoprotein B, is expressed after viral replication with γ_1_ late kinetics (Pellett et al., 1985). HSV1-Che is a recombinant strain derived from the bacterial artificial chromosome (BAC) pHSV1(17^+^)-Lox of the clinical isolate strain 17^+^ and, like all BAC-derived HSV-1 strains, lacks one of the three viral origins-of-replication (Nagel et al., 2008). As BAC-derived HSV-1 strains are impaired in some animal infection models, in particular in reactivation from latency (Sawtell and Thompson, 2014), we compared the infection of the BAC-derived HSV1(17^+^)Lox-Che and HSV1(17^+^)Lox strains to a low-passage preparation of HSV-1 strain 17^+^, their parental clinical isolate. The clinical isolate had replicated to slightly higher titers than HSV1(17^+^)Lox-Che or HSV1(17^+^)Lox in the skin explants (Fig. 3C).

**Figure 3.**
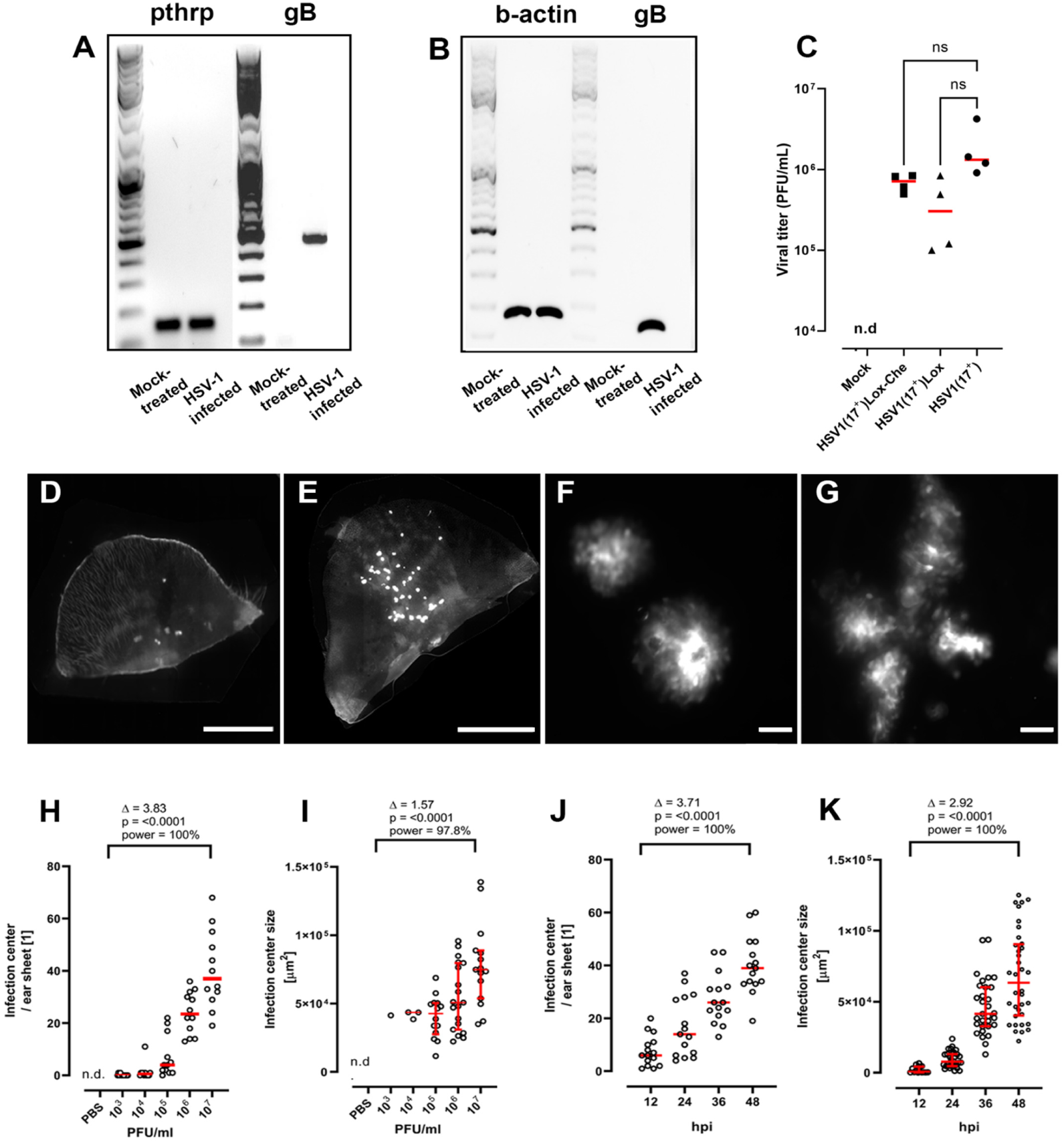
*Ex vivo* infection of murine skin sheets with HSV-1. Murine ear sheets were mock-treated or infected with HSV1(17^+^)Lox-Che (A-K), HSV1(17^+^)Lox (C), or HSV1(17^+^) (C) at 1 x 10^7^ (A-G, J, K) or at 10^3^ to 10^7^ (H, I) pfu/filter paper for 48 h (A-I) or 12 to 48 h (J, K). (A) At 48 hpi, the DNA was isolated from the explants, and the amount of murine genomes was determined by PCR for parathyroid hormone-related protein (pthrp) and the amount of HSV-1 genomes by PCR for UL27 encoded gB. (B) At 48 hpi, the RNA was isolated, and the amount of murine transcripts was determined by RT-PCR for β-actin, and the amount of HSV-1 transcripts by PCR for gB. (C) At 48 hpi, the amount of infectious HSV-1 was determined by plaque assay in 4 independent biological replicates. (D – G) Ear sheets inoculated after depilation (D, F, G) or after mock-treatment without depilatory cream (E) were fixed, and the infectious centers as indicated by the HSV-1 Che expression in whole-mounts were analyzed by fluorescence microscopy at low (D, E; scale bar 500 μm) or high magnification (F, G; scale bar 50 μm). (H – K) Ear sheets were infected with increasing amounts of HSV1-Che for 48 h (H, I) or at 1 x 10^7^ pfu/filter paper for 12 to 48 h (J, K), fixed, and the number (H, J) and the size (I, K) of the infection centers were determined. Mean (H-K; red) with standard-error of the mean (I, K; red), effect size (Δ), p-value, and post hoc statistical power are indicated from compiled data of 3 independent biological replicates.

**Table 1.**
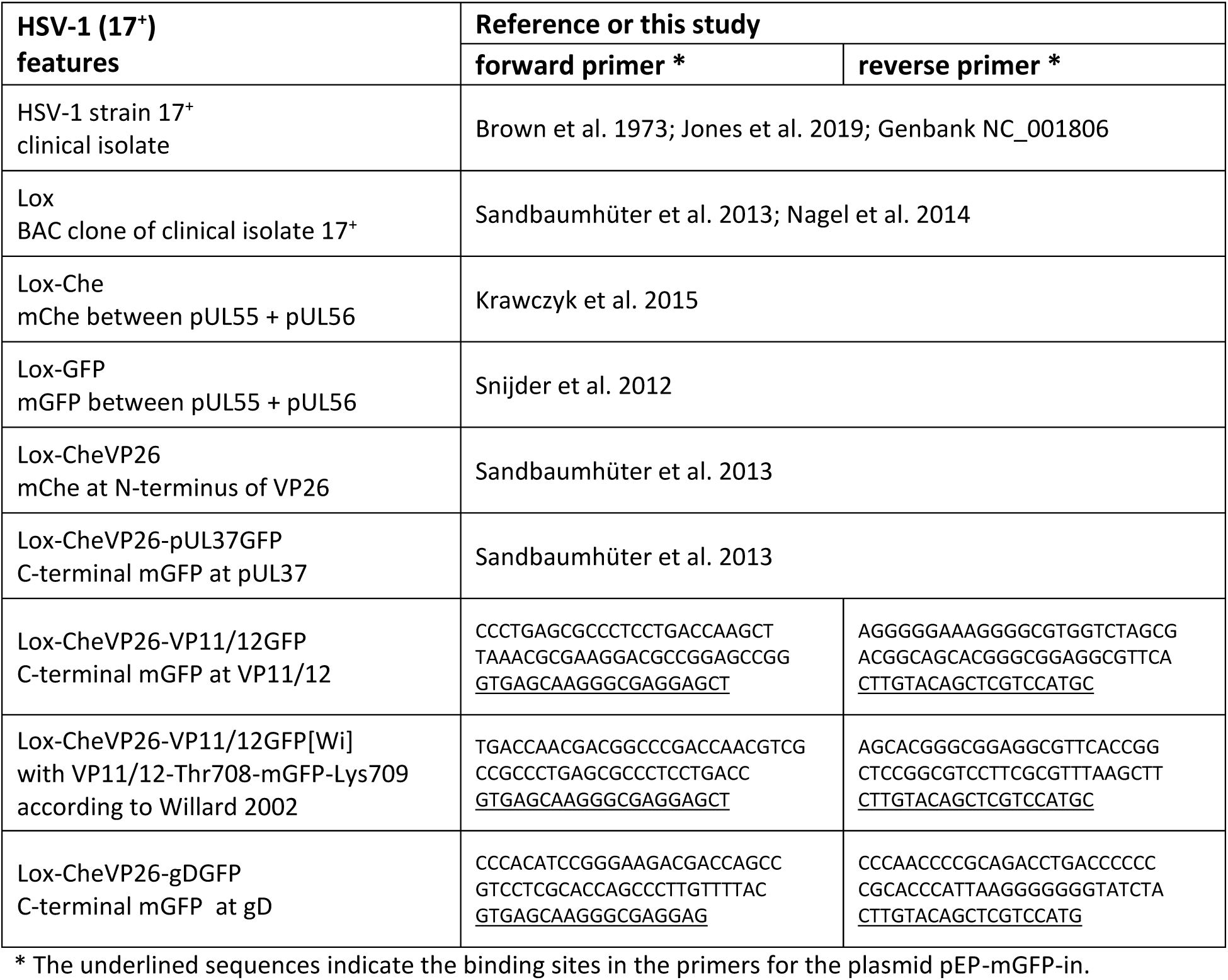
Published and generated HSV-1 strains used in this study.

Fluorescence microscopy of whole-mount explants revealed several infection centers expressing HSV-1-encoded mCherry in the regions that had been inoculated via the filter papers but not in the surrounding skin (Fig. 3D). Without depilation, HSV1-Che rarely established infection centers, and their numbers varied stochastically among explants (Fig. 3E). Following infection using the HSV-1-soaked filter papers, the size of the infection centers was heterogeneous, and a closer inspection at higher magnification suggested that some had originated from a single focus, and thus possibly from a single infected cell (Fig. 3F), while others contained multiple foci that had coalesced into a larger infected region (Fig. 3G). At a dose of 10^6^ PFU/filter or more, all skin explants contained infection centers (Fig. 3H). At 48 hpi, at a dose of 10^4^ to 10^6^ PFU/filter, the median sizes of the infection centers were rather similar but larger at 10^7^ PFU/filter (Fig. 3I). At 10^7^ PFU/filter, small fluorescent infection centers had formed already within 12 hpi, and their number (Fig. 3J) and size (Fig. 3K) increased over time.

With this novel protocol, we could robustly and reproducibly inoculate with a higher virus dose than by pipetting the inoculum directly onto the skin. The increasing number of infection centers, either with a higher dose or prolonged infection time, might indicate HSV-1 spread from primary infection centers and the formation of secondary infection centers or a heterogeneous expression of intrinsic antiviral proteins. In contrast to conventional plaque assays with cell monolayers (Grosche et al., 2019), we did not include any agents such as neutralizing antibodies, methylcellulose, or agarose. However, we assume that there was only lateral cell-to-cell in this air-liquid-interface culture. Our data are consistent with the notion that increasing the dose or the infection time augmented the likelihood of forming large coalescent infection centers derived from several small infection centers.

### HSV-1 spreads in the murine epidermis but not to the dermis

Next, we characterized the organization of the infection centers by 2-photon fluorescence microscopy. The infection centers were also detected with this technique as early as 12 hpi with HSV1-Che (Fig. 4A). Two-dimensional (Fig. 4B) and three-dimensional (Fig. 4C, 4D) reconstructions indicated that the infection centers were limited to the epidermis and had expanded more in the x-y than in the z directions. The volume of the infectious centers as measured from the rendered surface of HSV1-Che-expressing tissue (Fig. 4E, 4F) increased exponentially over time (Fig. 4G). Infection with HSV1-GFP, which expresses soluble monomeric GFP as a reporter, yielded centers of similar size with similar kinetics (Fig. 4H).

**Figure 4.**
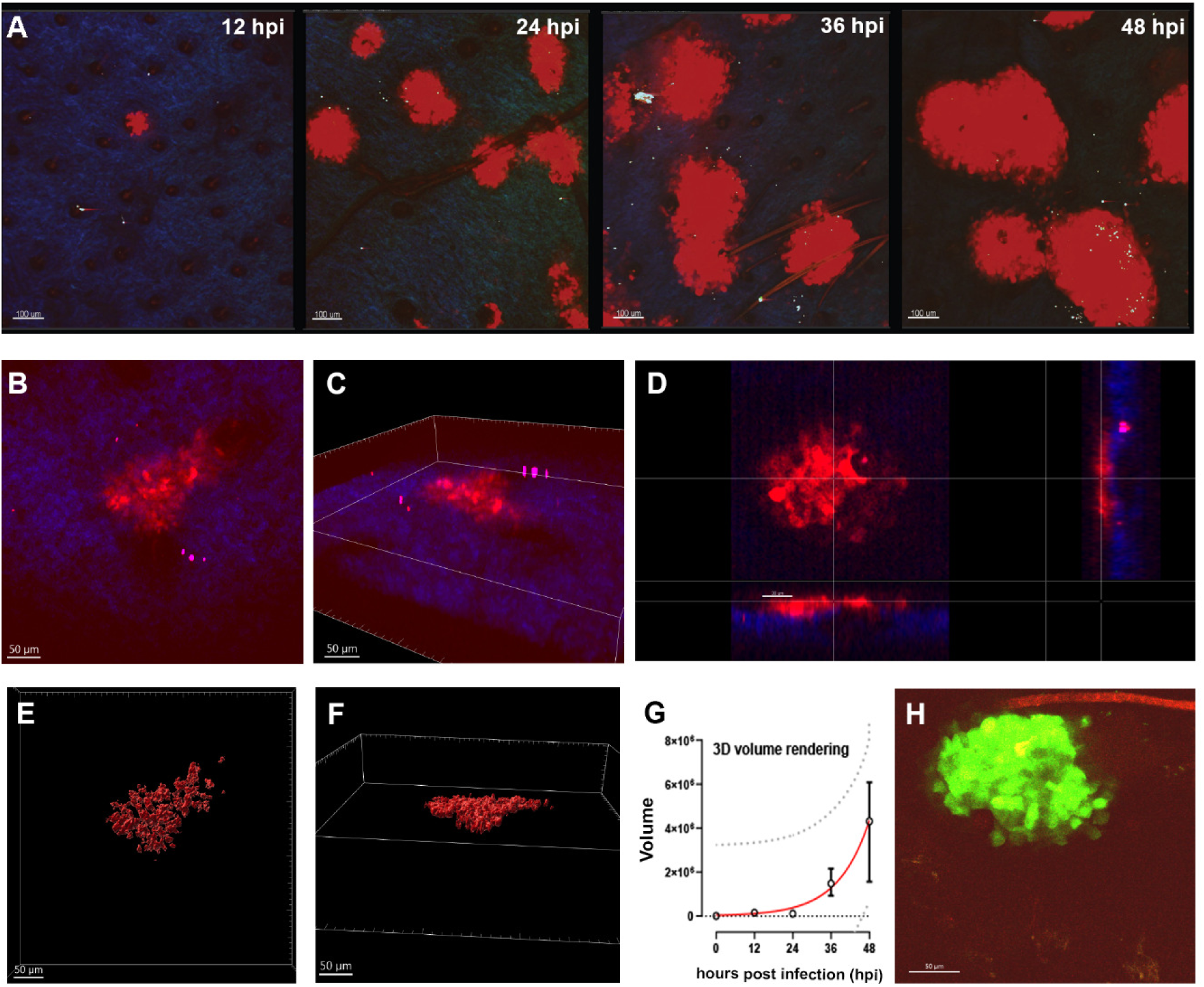
HSV-1 spreads in the epidermis but not the dermis of murine skin explants. (**A-H**) Ear sheets were infected with HSV1-Che or HSV1-GFP at 1 x 10^7^ pfu/filter paper, fixed, and analyzed by 2-photon microscopy for HSV1-Che (A – G; red) or HSV1-GFP (H; green) expression and with the second harmonic signal for collagen (A-D; blue). Expansion of HSV1-Che infection centers over time (A). Views of infectious centers in 2-dimensions (B), 3-dimensions (C-D), or with rendered nuclei (E-F) at 48 hpi. The volume of the infection centers was measured until 48 hpi based on the rendered nuclei from two independent experiments (G; median, IQR; red line, fitted exponential growth curve; 95% prediction bands, dotted line).

HSV1-Che infection led to moderate to marked multifocal epithelial hyperplasia and mild, multifocal, orthokeratotic hyperkeratosis, a thicker stratum corneum, with ballooning degeneration and multifocal intranuclear inclusion bodies within keratinocytes as indicated by HE staining (Fig. 5A) and immunolabeling for pan-cytokeratin (Fig. 5B). The HSV-1 single-strand DNA binding protein ICP8, which is essential for the nuclear HSV-1 DNA replication (Weller et al., 1983), was localized exclusively in the nuclei of keratinocytes but were not present in cells of the underlying dermis (Fig. 5C). Hence, HSV-1 infection was limited to the epidermis and did not spread to the dermis within 48 hpi in the murine skin explant cultures. In non-infected skin explants, there was a mild, multifocal to coalescing epithelial hyperplasia accompanied by mild, multifocal to coalescing, orthokeratotic hyperkeratosis (Fig. 5D, E), but the antibody directed against HSV-1 ICP8 did not react with the uninfected control (Fig. 5F). The mild hyperplasia and mild hyperkeratosis of the uninfected skin explants was likely due to a reaction to the depilatory creme.

**Figure 5.**
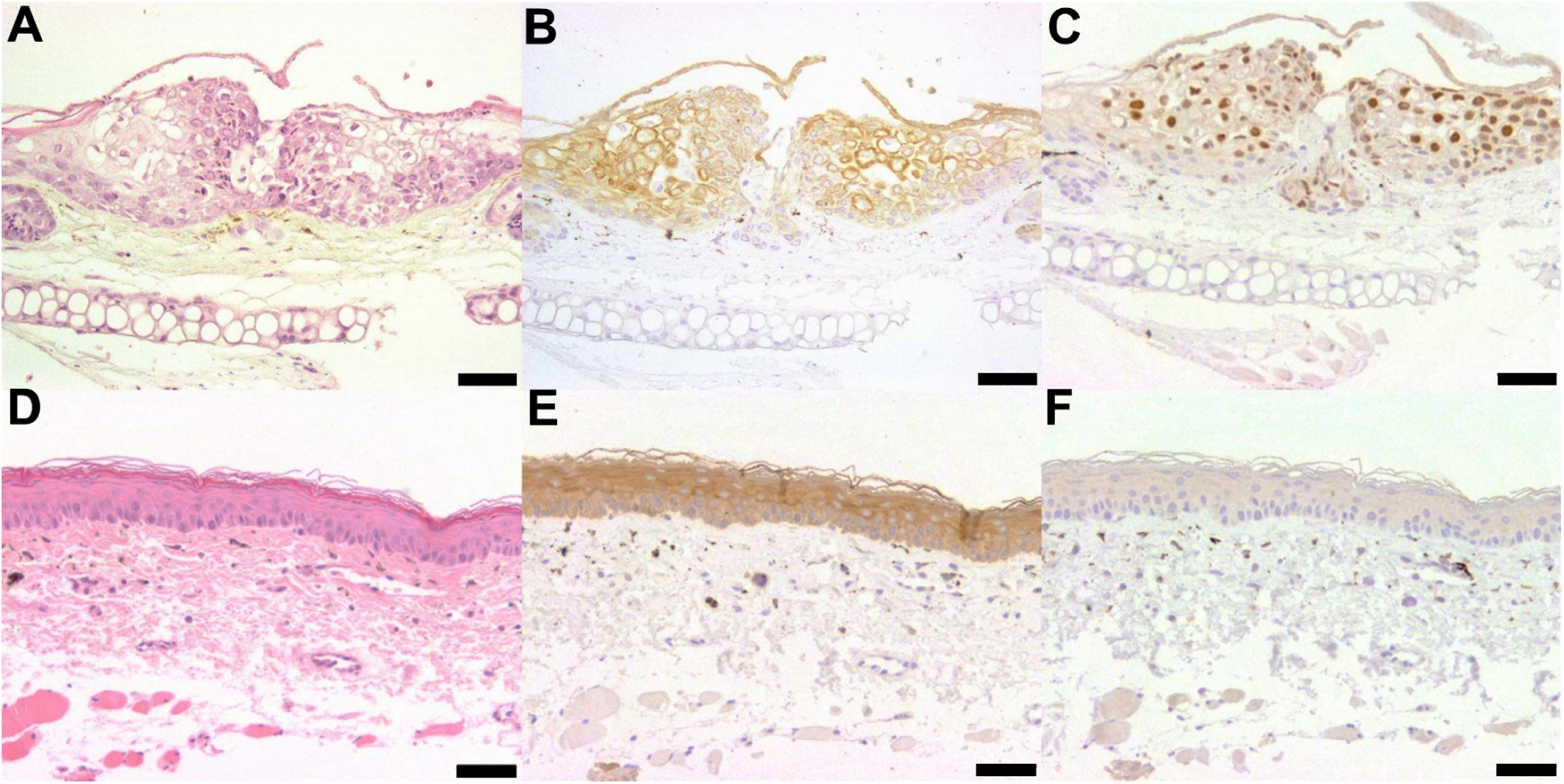
HSV-1 induces epidermal hyperplasia in murine skin. Skin explants were infected with HSV1-Che at 1 x 10^7^ pfu/filter paper (A,B,C) or mock infected (D,E,F), stained with hematoxylin and eosin (A,D) and labeled for pan-cytokeratin (B; E) or the HSV-1 single-strand DNA binding protein ICP8 (C,F). Scale bar 50 μm.

### Characterization of dual-color HSV-1 strains

To be able to monitor the different stages of the infection cycle, and to characterize the intracellular distribution of different HSV-1 assembly intermediates, we used strains in which the small capsid protein VP26 had been tagged at its N-terminus with mCherry (Buch et al., 2017; Döhner et al., 2018; Ivanova et al., 2016; Sandbaumhüter et al., 2013), and the inner tegument protein pUL37 (Sandbaumhüter et al., 2013), the outer tegument protein VP11/12 (Lin et al., 2013; this study), or the envelope protein gD (this study) with GFP (Table 1). Restriction digest analyses of the genomes of the novel reporter strains showed that HSV1-CheVP26-VP11/12GFP with GFP added directly to its C-terminus (not shown), HSV1-CheVP26-VP11/12GFP[Wi] with GFP inserted 10 residues upstream of its C-terminus as reported before (Willard, 2002), and HSV1-CheVP26-gDGFP with GFP at its C-terminus showed the expected band shifts in comparison to the parental strains HSV1(17^+^)Lox or HSV1-CheVP26 upon digestion with BamHI (Fig. 6A) or SalI (Fig. 6A). Tagging HSV1-VP26 (Fig. 6B) as reported before (Sandbaumhüter et al., 2013), or tagging HSV1-VP11/12 (Fig. 6C) did not impair replication in single-step growth curves, whereas tagging HSV1-gD delayed the release of infectious virus from the infected Vero cells (Fig. 6D). Accordingly, the average plaque sizes of the HSV-1 strains with a GFP tag on pUL37 or VP11/12 were similar to that of the parental strain, while tagging gD had reduced the plaque size considerably (Fig. 6E).

**Figure 6.**
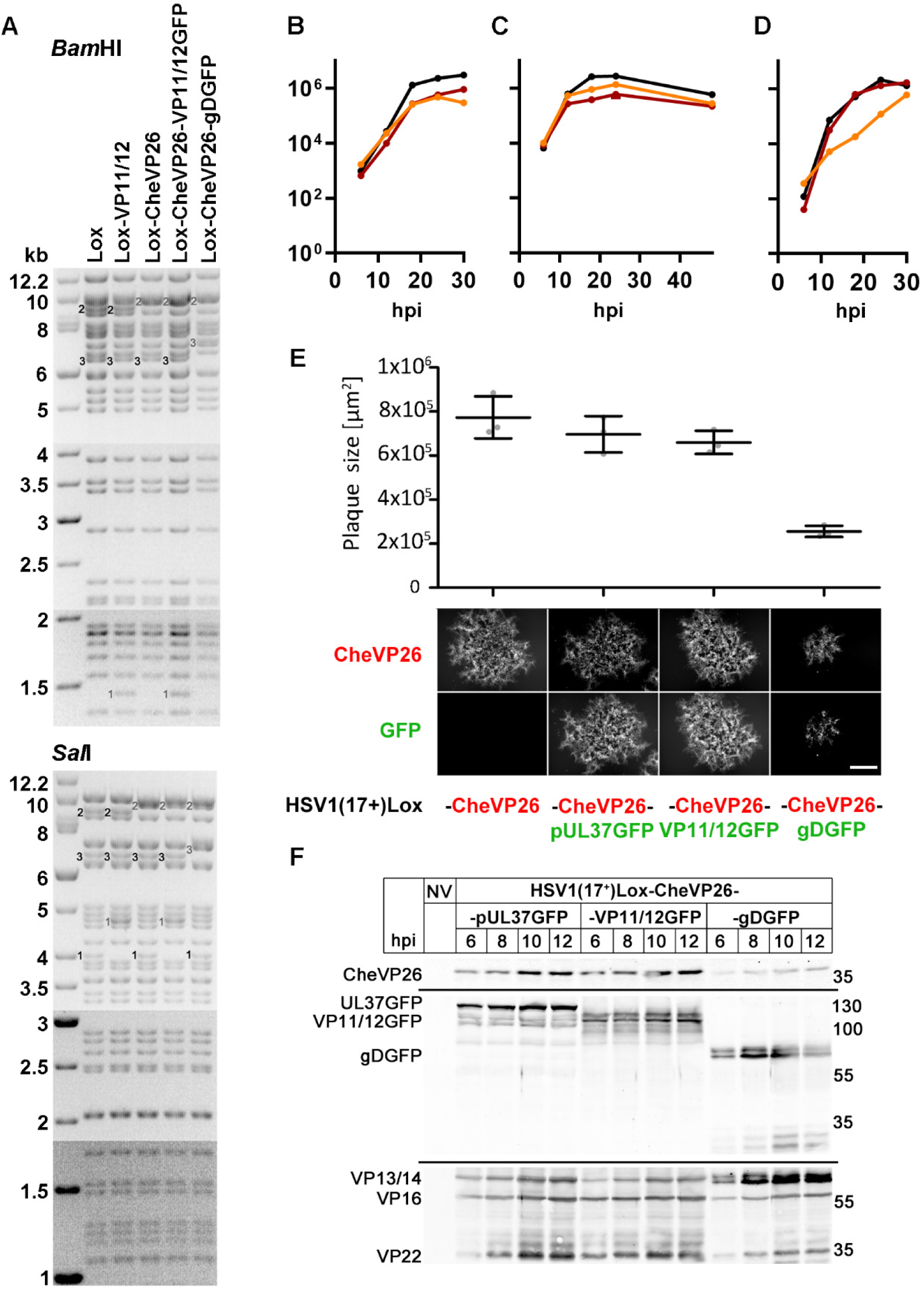
Characterization of the dual-color HSV-1 strains. (A) Agarose gel of genome restriction digests with *BamH*1 or Sal1 of HSV1(17^+^)Lox, Lox-VP11/12GFP, Lox-CheVP26, Lox-CheVP26-VP11/12GFP[Wi], and Lox-CheVP26-gDGFP. The fragments had shifted as expected due to adding mGFP to VP11/12, mCherry to VP26, or mGFP to gD. The sizes of the molecular weight marker bands are indicated in kB. The black numbers indicate the parental forms and grey numbers the tagged forms. (B-D) For single-step growth curves, Vero cells were infected with HSV1(17^+^)Lox (black in B-D), Lox-CheVP26 (red in B-D), Lox-CheVP26-pUL37GFP (orange in B), Lox-CheVP26-VP11/12GFP[Wi] (orange in C), or Lox-CheVP26-gDGFP (orange in D) at 5 pfu/ cell, and infectious virus released from infected cells into the medium at the indicated time points was titrated on Vero cells. (E) To determine plaque sizes, Vero cells were infected with HSV1-CheVP26, –CheVP26-UL37GFP, –CheVP26-VP11/12GFP[Wi], or –CheVP26-gDGFP in a dilution series, and at 2 dpi Cherry-or GFP-positive plaques were documented by live-cell imaging, and the plaques sizes were measured using FIJI. (F) Immunoblot. Vero cells were mock-treated (NV) or infected for indicated times at 10 pfu/cell with HSV1(17^+^)Lox-CheVP26-pUL37GFP, Lox-CheVP26-VP11/12GFP[Wi], or Lox-CheVP26-gDGFP, and cell lysates were analysed by immunoblot with antibodies raised against RFP, GFP, or various HSV-1 antigens (Remus V).

Upon infection of permissive Vero cells at a high multiplicity, all dual-color HSV-1 strains progressed rather synchronously through virion morphogenesis, although HSV1-CheVP26-gDGFP was a bit delayed consistent with its slower plaque expansion (c.f. Fig. 6E). All cytoplasmic but not nuclear CheVP26 capsids (Fig. 7Ai, Bi, red in Aiv, Av, Biv, and Bv) co-localized with inner tegument protein pUL37GFP (Fig. 7Aii, Bii green in Aiv, Biv) as reported before (Sandbaumhüter et al., 2013), while the transcriptional activators and tegument proteins HSV1-VP16 (Fig. 7Aiii, blue in Av) and VP22 (Fig. 7Biii, blue in Bv) were diffusively distributed in the nucleus and in large cytoplasmic accumulations that also contained cytoplasmic capsids, but not present on all cytoplasmic capsids. In contrast, many but not all cytoplasmic capsids (Fig. 7Ci, Di, red in Civ, Cv, Div and Dv) co-localized with outer tegument protein VP11/12GFP (Fig. 7Cii, Dii green in Civ, Div). Moreover, VP11/12GFP also localized to nuclear domains in close proximity to nuclear VP26 structures (Fig. 7Cii, Dii, green in Civ, Div) and to the nuclear rim (not shown). Moreover, many but not all cytoplasmic capsids colocalized (Fig. 7Ei, Fi red in Eiv, Ev, Fiv and Fv) with the envelope protein gDGFP (Fig. 7Eii, Fii green in Eiv and Fiv). These data suggest that the HSV-1 dual-color reporter viruses faithfully indicated the sequential acquisition of tegument proteins to capsids (reviewed in Döhner et al., 2024). While the inner tegument protein pUL37GFP could be detected on all cytoplasmic CheVP26 capsids, capsids colocalizing with the outer tegument proteins VP16 (Fig. 7Aiii, Ciii, Eiii; blue in Av, Cv, Ev) and VP22 (Fig. 7Biii, Diii, Fiii; blue in Bv, Dv, Fv) as well as the envelope protein gDGFP were likely in the process of secondary capsid envelopment.

**Figure 7:**
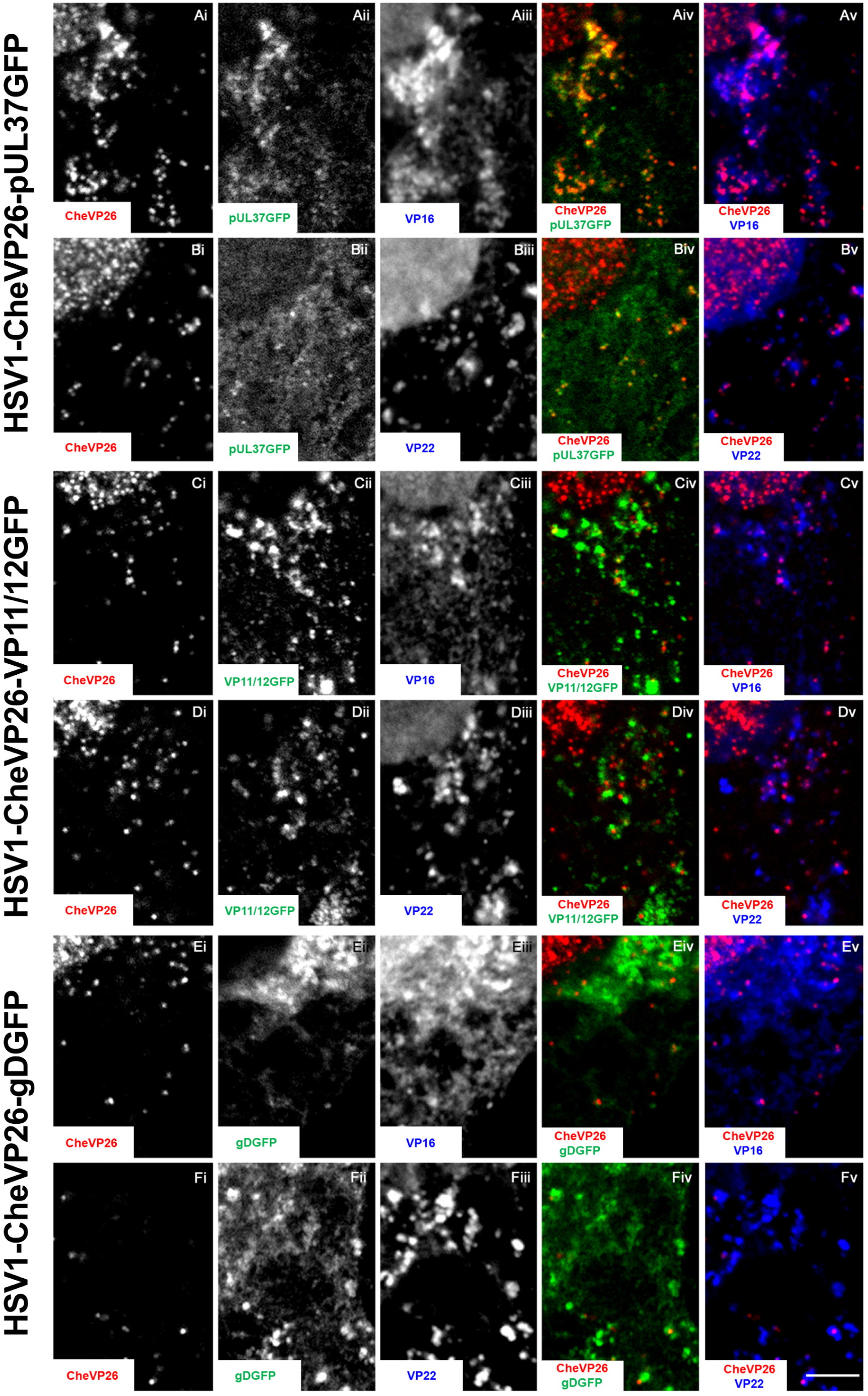
HSV-1 dual color strains for tracking different assembly intermediates. Vero cells were infected at 10 pfu/cell (3.75 x 10^6^ pfu/mL) with HSV1-CheVP26-pUL37GFP for 10 h (A,B), –CheVP26-VP11/12GFP[Wi], for 8 h (C,D), or –CheVP26-gDGFP for 12 h (E,F). The cells were fixed with 3% paraformaldehyde/PBS, permeabilized with 0.1% TX-100, labelled, and analyzed for HSV-1 expression of the capsid protein CheVP26 (i; red in iv and v), the inner tegument protein pUL37GFP (Aii, Bii; green in Aiv, Biv), the outer tegument protein VP11/12GFP (Cii, Dii; green in Civ, Div), the envelope protein gDGFP (Eii, Fii; green in Eiv, Fiv), VP16 (mAb LP1; Aiii, Ciii, Eiii, blue in Av, Cv, Ev), and VP22 (mAb AGV30; Biii, Diii, Fiii, blue in Bv, Dv, Fv) by confocal fluorescence microscopy. Scale bar 5 µm.

### HSV-1 capsid assembly and secondary envelopment in keratinocytes

Next, we infected skin explants with HSV1-CheVP26 to stage the infection in individual cells, and to monitor the formation of intracellular assembly intermediates. Confocal fluorescence microscopy of sections through infection centers cut perpendicular to the skin surface shows that the host chromatin was confined to the nuclear periphery in infected cells when compared to the surrounding uninfected cells (DAPI; Fig. 8Ai, Ci, Di, Ei). Instead, the nuclei contained large accumulations of CheVP26 capsids; moreover, numerous individual CheVP26 particles were detected mostly in the nucleoplasm, but also in the cytoplasm (Fig. 8Aii, Cii, Dii, Eii). Such marginalization of host chromatin also occurs during HSV-1 infection of cultured cells (Aho et al., 2021; Bosse et al., 2015). Labeling for nuclear pore proteins indicated that the nuclear envelopes also contained capsids (green in Fig. 8Bii). Infection with HSV1-CheVP26-pUL37GFP showed co-localization of cytoplasmic but not nuclear CheVP26 capsids with the inner tegument protein pUL37GFP (Fig. 8Civ). Similarly, after infection with HSV1-CheVP26-VP11/12GFP or HSV1-CheVP26-gDGFP, cytoplasmic but not nuclear CheVP26 capsids colocalized with the outer tegument protein VP11/12GFP (Fig. 8Div) or with the envelope glycoprotein gDGFP (Fig. 8Eiv), respectively.

**Figure 8.**
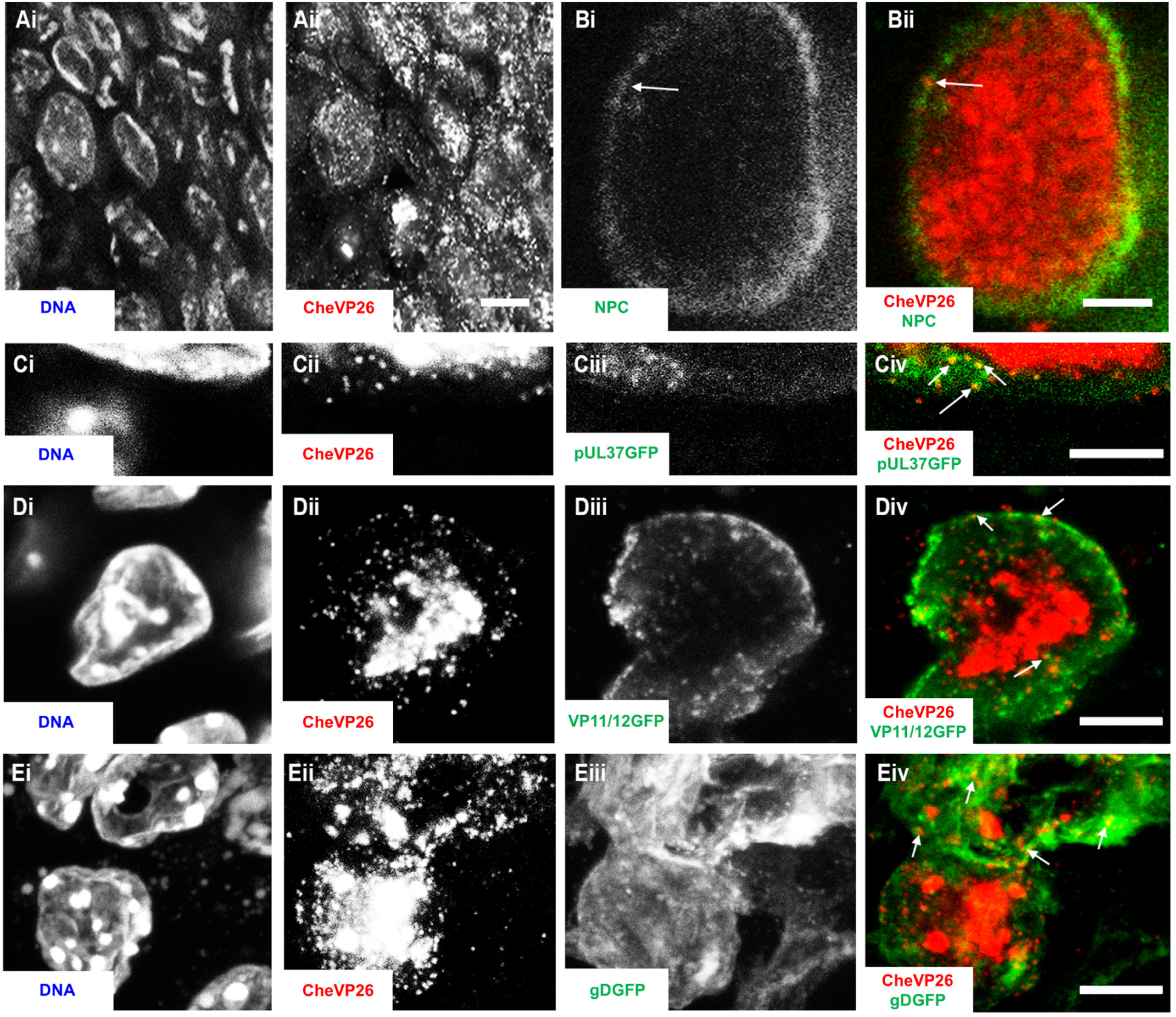
HSV-1 assembly and chromatin marginalization in murine skin keratinocytes. Ear sheets were infected with HSV1-CheVP26 (A, B), –CheVP26-pUL37GFP (C), –CheVP26-VP11/12GFP[Wi] (D), or –CheVP26-gDGFP (E) at 1 x 10^7^ pfu/filter paper for 48 hpi. Cryosections were stained for DNA (DAPI; Ai, Ci, Di, Ei), in some cases labeled for the nuclear pores (Bi; green in Bii), and analyzed for expression of the HSV-1 capsid protein CheVP26 (Aii, Cii, Dii, Eii; red in Bii, Civ, Div, Eiv), the inner tegument protein pUL37GFP (Ciii; green in Civ), the outer tegument protein VP11/12GFP (Diii; green in Div), or the envelope protein gDGFP (Eiii; green in Eiv) by confocal fluorescence microscopy. Scale bar: 25 µm (A) or 5 µm (B-E).

Next, we analyzed the HSV-1 infection of murine skin explants by transmission electron microscopy of sections cut perpendicular through the infection centers identified by their Che fluorescence before the processing for electron microscopy. The depilation had removed some but not all cells of the apical stratum corneum as indicated by a remaining prominent dark keratin layer (Fig. 9A). We detected HSV1 capsids predominantly in the nuclei, and less frequent in the cytoplasm of keratinocytes in the stratum granulosum (Fig. 9C), the stratum spinosum (Fig. 9C, D), and the stratum basale (Fig 9E).

**Figure 9.**
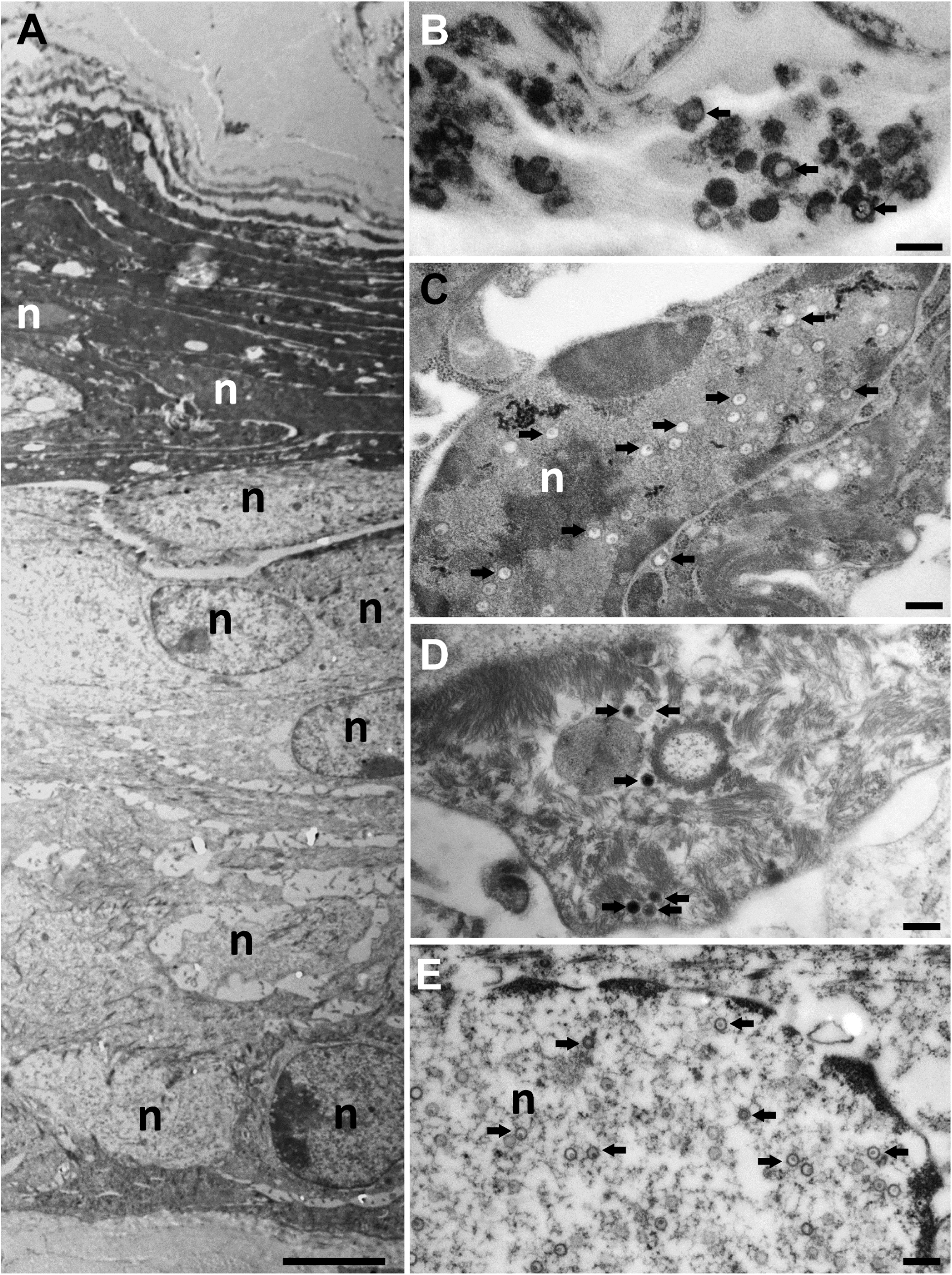
Ultrastructure of the HSV-1 infected epidermis in murine skin explants. Ear sheets were infected with HSV1-Che at 1 x 10^7^ pfu/filter paper for 48 hpi, fixed, embedded in resin, cut in ultrathin sections perpendicular to the skin surface, and analysed by electron microscopy. (A) An overview at low magnification shows the four layers of the epidermis; with keratinocytes from the stratum corneum at the apical surface, stratum granulosum, and stratum spinosum to the stratum basale at the bottom of the image. (B-D) At higher magnifications, viral capsids (black arrows) can be recognized in all layers of the epidermis, extracellular viral structures in the stratum corneum (B), capsids in the nuclei, and the cytoplasm of keratinocytes in the stratum granulosum (C), the stratum spinosum (D), and the stratum basale (E). Scale bar: 2 µm in A, 200 nm in B-E; nuclei are labeled with n.

In the nuclei of these cells HSV-1 capsid assembly intermediates were found such as empty A capsids (Fig. 10A, white arrows), B capsids with an internal protein core (outlined white arrows), and C capsids containing viral genomes (black arrows). Furthermore, we detected all known HSV1 egress stages which are primary virions in the perinuclear space (Fig. 10B), cytosolic capsids (Fig 10C), capsids in the process of secondary envelopment (Fig. 10D, E), and intracellular vesicles harboring apparently intact virions (Fig. 10F). In the stratum corneum, containing dead keratinocytes, extracellular virus structures were found (Fig 9B). These results indicate efficient nuclear capsid egress and virion assembly. In summary, the fluorescence and electron microscopy data indicate that HSV-1 had productively infected the keratinocytes, that all HSV-1 assembly intermediates had been formed, and that cytosolic capsids had recruited the inner tegument protein pUL37GFP and the outer tegument protein VP11/12GFP and undergone secondary envelopment on cytoplasmic membranes containing gDGFP.

**Figure 10.**
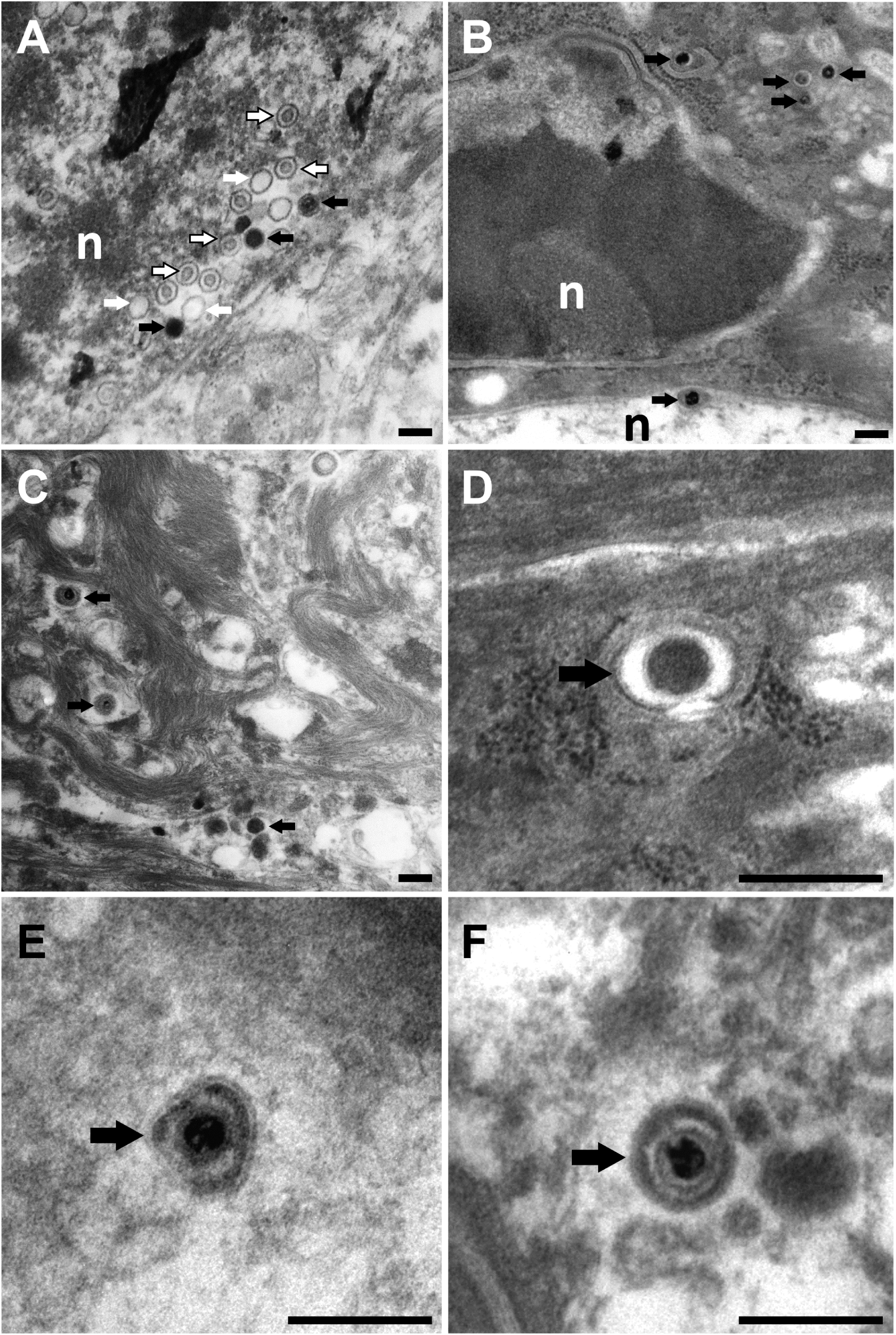
HSV-1 capsid assembly, secondary envelopment and virion production in keratinocytes. Ear sheets were infected with HSV1-Che at 1 x 10^7^ pfu/filter paper for 48 hpi and processed for electron microscopy. The infected cells contain all HSV-1 assembly intermediates, namely A capsids (white arrows in A), B capsids (outlined arrows in A), C capsids (black arrows in A), primary enveloped virions in the lumen between inner and outer nuclear membrane (black arrows in B), cytosolic capsids (black arrows in C), capsids in the process of secondary envelopment (black arrow in D and E), and intracellular vesicles harboring apparently intact virions (black arrow in F). Scale bar 200 nm; nuclei are labeled with n.

## DISCUSSION

Intact skin and mucosal barriers are essential for survival, but many viruses replicate in barrier tissues, and local immune responses often fail to provide sterilizing immunity. To better understand virus spread within barrier tissues, we established a novel murine infection model to directly observe virus-infected cells within the first few hours of infection. We inoculated from the apical “outside” to investigate the very early events of HSV-1 infection before the onset of clinical symptoms and without the induction of prior inflammation. HSV-1 and HSV-2 can cause painful skin lesions such as *herpetic whitlow* (“whitlow finger”) in health care workers (Feder, 1983), outbreaks of *herpes gladiatorum* (“mat herpes”) in contact-sports athletes (De Bernardo, 1992), early-onset infection after cosmetic tattooing (Serup, 2023), and *eczema herpeticum* in a subgroup of patients with atopic dermatitis (Traidl et al., 2023).

Although many small mammals can be infected with HSV-1 via several routes, murine models are favored due to the huge variety of host-specific markers and transgenic knock-in and knock-out strains (Hill and Shimeld, 1998; Hussain et al., 2024; Kollias et al., 2015). While many studies focus on the skin of the flank or the footpad (Allan et al., 2003; Bedoui et al., 2009; Gebhardt et al., 2011; Mackay et al., 2012; Wakim et al., 2008), we infected the ear skin of six-to nine-week-old mice because these tissue samples are highly suited for different imaging modalities. In the long run, this approach is also amenable to multi-photon fluorescence microscopy analyses of cutaneous HSV-1 spread in living *ex vivo* tissues and even in animals. Infection of the ear pinnae of several murine strains with different HSV-1 strains mimics many aspects of human pathogenesis, as it leads to acute skin lesions, the infiltration of immune cells, the establishment of latency in sensory neurons of the cervical ganglia, and zosteriform spread back to the skin (Harbour et al., 1981; Hill et al., 1975; Proença et al., 2016; Puttur et al., 2010).

Instead of the commonly used skin scarification by sandpaper, needling, tattooing, or tape stripping, all of which can induce innate immune responses before infection (Amberg et al., 2017; Reichert et al., 2023), we used a short depilation treatment to allow HSV-1 access to living apical keratinocytes while leaving the dermal-epidermal junction intact. A similar strategy using tape stripping or depilatory cream has been used to investigate HSV-1 infection of gamma-delta T cells and Langerhans cells in murine ear explant cultures (Puttur et al., 2010). The thioglycolate acids in depilatory creams reduce inter– and intra-protein disulfide bonds; hairs break off above their roots and are removed along with the intercellular lipids and superficial dead keratinocytes, the corneocytes, rich in keratin, filaggrin, loricrin, and other proteins (Lee et al., 2008). We used high-titer HSV-1 inocula of high quality with low particle/pfu ratios (Grosche et al., 2019), and limited their application to the areas below the filter papers.

Although the air-liquid-interface skin cultures were fed only with culture medium and no longer by the blood circulation, we could detect single HSV-1 infected keratinocyte as early as 12 hpi and monitor the cutaneous spread and the development of infectious centers for up to 96 hpi using reporter strains expressing fluorescent proteins. The increasing size of the infection centers over time indicates HSV-1 cell-to-cell spread (c.f. Fig. 3K), but their increasing number suggests the formation of additional secondary infection centers (c.f. Fig. 3G; 3J). Similar to murine epidermal sheets but not cornified skin from the tails (Petermann et al., 2015; Rahn et al. 2015; Rahn et al., 2017), the ear skin was also more susceptible to primary HSV-1 infection from the outside after the corneocytes had been removed by depilation. However, Rahn *et al*. (2017) could not infect keratinocytes of the tail skin explants even after wounding or removal of the apical stratum corneum. While it is difficult to estimate the amount of infectious HSV-1 units that had been released from the filter papers to the skin surface in our protocol, in Rahn et al. (2017), the multiplicity of infection might have been lower. In addition, their inocula might have contained more defective viral particles or cellular debris that via stimulation of pattern-recognition receptors might have induced an antiviral state in the skin. Combining the use of inocula of high titer and of high quality with their application after depilation and to a defined surface area via filter papers, which prevented their dilution with the culture medium, enabled us to obtain robust and reproducible numbers of infection centers among different experiments.

Our novel *ex vivo* culture of terminally differentiated skin is suitable to characterize the phenotype of HSV-1 mutants impaired in evading innate immune responses, as well as skin infection with varicella-zoster virus, poxvirus, papillomavirus, and others. Furthermore, it should be feasible to infect the ear pinnae of animals rather via the small pieces of filter paper to deliver a rather high dose of viral inocula locally, and to study the contribution of locally-induced interferons and cytokines, infiltrating immune cells, and the neuro-immune crosstalk to skin lesion formation and healing. Moreover, such *ex vivo* and *in vivo* skin infection could be used to investigate the potency of potential novel anti-viral small chemical compounds. As pioneered by Knebel-Mörsdorf and her team (De La Cruz et al., 2021; Möckel et al., 2022), we envision to characterize HSV-1 infection of skin explants from atopic dermatitis patients with defined candidate single nucleotide polymorphisms that are hypothesized to correlate with their susceptibility to eczema herpeticum in atopic dermatitis. We also have identified RNAse7, the short-form of thymic stromal lymphopoietin and collagen XXIIIα as host proteins that contribute to the susceptibility to *eczema herpeticum* in patients with atopic dermatitis, and that modulate HSV-1 infection in human keratinocytes (Kopfnagel et al., 2020; Zeitvogel et al., 2024; Chopra et al., 2024).

## MATERIALS AND METHODS

### Cells and antibodies

BHK-21 (ATCC CCL-10) and Vero cells (ATCC CCL-81) were cultured at 37°C and 5% CO_2_ in MEM supplemented with 1% non-essential amino acids (Cytogen, Wetzlar, Germany) and 10% or 7.5% fetal bovine serum (Good Forte; PAN-Biotech, Aidenbach, Germany) and passaged twice per week. In the plaque assays, we used human antibodies including HSV-1 neutralizing antibodies to limit plaque formation to cell-to-cell spread (Beriglobin; CSL Behring, Hattersheim am Main, Germany; (Grosche et al., 2019). For the immunofluorescence microscopy experiments, we used primary antibodies directed against host or HSV-1 proteins, and secondary antibodies cross-adsorbed for species specificity and attached to fluorescent dyes (Table 2). To competitively block binding of any primary or secondary antibodies to the HSV-1 encoded Fc receptor of glycoproteins gE/gI, we used human sera of HSV-1 seronegative volunteers (Döhner et al., 2006; Grosche et al., 2019)

**Table 2.**
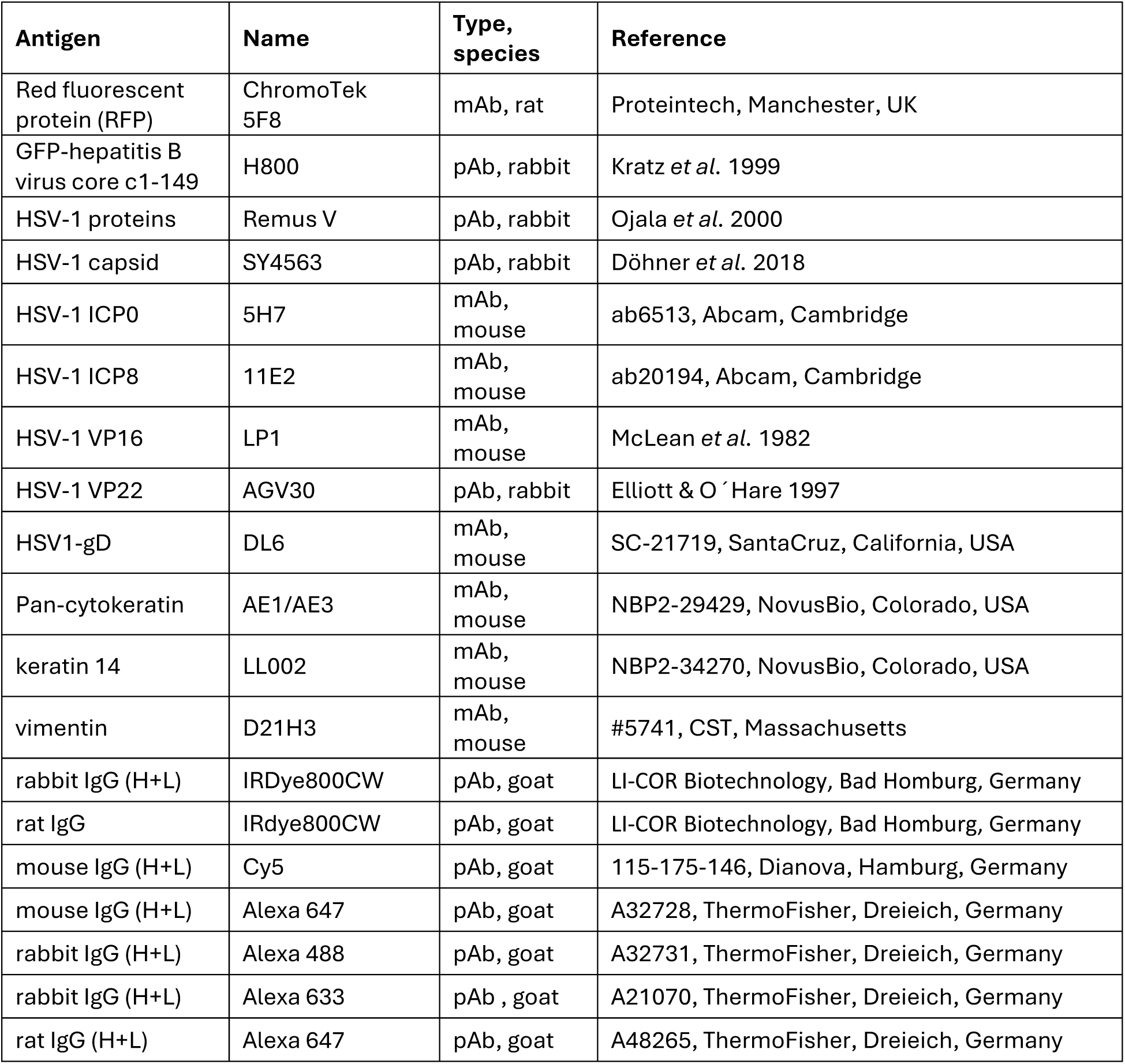
Primary and Secondary antibodies.

| Antigen | Name | Type, species | Reference |
| --- | --- | --- | --- |
| Red fluorescent protein (RFP) | ChromoTek 5F8 | mAb, rat | Proteintech, Manchester, UK |
| GFP-hepatitis B virus core c1-149 | H800 | pAb, rabbit | Kratz *et al.* 1999 |
| HSV-1 proteins | Remus V | pAb, rabbit | Ojala *et al.* 2000 |
| HSV-1 capsid | SY4563 | pAb, rabbit | Döhner *et al.* 2018 |
| HSV-1 ICP0 | 5H7 | mAb, mouse | ab6513, Abcam, Cambridge |
| HSV-1 ICP8 | 11E2 | mAb, mouse | ab20194, Abcam, Cambridge |
| HSV-1 VP16 | LP1 | mAb, mouse | McLean *et al.* 1982 |
| HSV-1 VP22 | AGV30 | pAb, rabbit | Elliott & O 'Hare 1997 |
| HSV1-gD | DL6 | mAb, mouse | SC-21719, SantaCruz, California, USA |
| Pan-cytokeratin | AE1/AE3 | mAb, mouse | NBP2-29429, NovusBio, Colorado, USA |
| keratin 14 | LL002 | mAb, mouse | NBP2-34270, NovusBio, Colorado, USA |
| vimentin | D21H3 | mAb, mouse | #5741, CST, Massachusetts |
| rabbit IgG (H+L) | IRDye800CW | pAb, goat | LI-COR Biotechnology, Bad Homburg, Germany |
| rat IgG | IRDye800CW | pAb, goat | LI-COR Biotechnology, Bad Homburg, Germany |
| mouse IgG (H+L) | Cy5 | pAb, goat | 115-175-146, Dianova, Hamburg, Germany |
| mouse IgG (H+L) | Alexa 647 | pAb, goat | A32728, ThermoFisher, Dreieich, Germany |
| rabbit IgG (H+L) | Alexa 488 | pAb, goat | A32731, ThermoFisher, Dreieich, Germany |
| rabbit IgG (H+L) | Alexa 633 | pAb, goat | A21070, ThermoFisher, Dreieich, Germany |
| rat IgG (H+L) | Alexa 647 | pAb, goat | A48265, ThermoFisher, Dreieich, Germany |

### Viruses and BAC mutagenesis

We used the clinical isolate HSV-1 strain 17^+^ (Brown et al., 1973; Jones et al., 2019) and the derived BAC strains HSV1(17^+^)Lox (HSV1-Lox for short; Nagel et al. 2014; Sandbaumhüter et al., 2013), HSV1(17^+^)Lox-Che (HSV1-Che; Krawczyk et al., 2015), and HSV1(17^+^)Lox-GFP (HSV1-GFP; Snijder et al., 2012; Table 1). Furthermore, we used HSV1(17^+^)Lox-CheVP26 with an monomeric Cherry tag on the N-terminus of the small capsid protein VP26 (HSV1-CheVP26; Sandbaumhüter et al., 2013), HSV1(17^+^)Lox-CheVP26-UL37GFP with additionally a monomeric GFP tag on the C-terminus of inner tegument protein pUL37 (HSV1-CheVP26-UL37GFP, Sandbaumhüter et al., 2013), with monomeric GFP on the C-terminus of the outer tegument protein VP11/12 (HSV1-CheVP26-VP11/12GFP; this study), with monomeric GFP inserted into the C-terminal third of VP11/12 between threonine 708 and lysine 709 at 10 residues upstream of the STOP codon as reported before (Willard, 2002; HSV1-CheVP26-VP11/12GFP[Wi]; this study), and HSV1(17^+^)Lox-CheVP26-gDGFP with monomeric GFP on the C-terminus of envelope protein gD (HSV1-CheVP26-gDGFP; this study). To generate the novel dual-color HSV1(17^+^)Lox strains for this study, we amplified PCR-fragments for mGFP-tagging from the plasmid pEP-mGFP-in (Sandbaumhüter et al., 2013) using the appropriate forward and reverse primers (Table 1), and used established methods for En passant BAC-mutagenesis (Bodda et al., 2020; Ivanova et al., 2016; Nagel et al., 2014; Tischer et al., 2006). After transformation into *E.coli* GS1783 carrying the parental BAC pHSV1(17^+^)Lox-CheVP26 and 2 rounds of RED-mediated recombination, the integrity of the BAC genomes was evaluated by restriction digest with BamH1, Sal1 (Fig. 5), Asc1, EcoR1, EcoRV, Not1, and Xho1 (not shown). The newly generated dual-color reporter viruses were reconstituted by transfecting the BAC DNA into Vero cells as described before (Sandbaumhüter et al., 2013). The insertion of coding sequences for mGFP was confirmed after reconstituting the respective HSV1(17^+^)Lox strains by sequencing 500 bp up and downstream of the respective insertion site (not shown).

Extracellular particles secreted from BHK-21 cells infected with HSV-1 at about 3 x 10^4^ PFU/mL (MOI of 0.01 PFU/cell) for 2 to 3 days until the cells had detached from the culture flasks were harvested by ultracentrifugation. The pellets were resuspended in MNT buffer (30 mM MES, 100 mM NaCl, 20 mM Tris, pH 7.4), aliquoted, snap-frozen in liquid N_2_, and stored at –80 °C in single-use aliquots as reported before (Grosche et al., 2019; Sodeik et al., 1997). The stocks had a titer of 2 x 10^8^ to 2 x 10^10^ PFU/mL on Vero cells and a genome/PFU ratio of 20 to 50 indicating very high-quality inocula with a low amount of defective, interfering particles (Döhner et al., 2006; Grosche et al., 2019).

### Plaque assay

The HSV-1 stocks as well as the explant-associated virus were titrated on Vero cells as reported before (Döhner et al., 2006; Grosche et al., 2019). The explants were removed from the filter papers, cut into small pieces of about 1 mm^3^, sheared (TissueRuptor II; Qiagen, Hilden, Germany) in MNT buffer, and freeze-thawed 3 times to release any cell-associated viruses. After the inoculation for 1 h, the cells were incubated in a regular culture medium containing 400 μg/mL human antibodies to limit the spread of extracellular virions (Beriglobin; CSL Behring, Hattersheim am Main, Germany).

### Infections

C57BL/6 mice were bred at the Central Animal Facility (MHH, Hannover Medical School). Female mice aged 6 to 14 weeks were sacrificed under anesthesia by inhalation of CO_2_ followed by cervical dislocation according to local animal welfare regulations, the ears were cut off, washed with 70% ethanol, treated with depilatory cream for 3 min (Veet PURE – hair removal crème for sensitive skin; Reckitt Benckiser Deutschland GmbH, Heidelberg, Germany), and the hairs and the cream were gently removed with cotton buds and PBS. S-shaped tweezers were used to separate the ears into apical and basal sheets. The sheets were glued with tissue adhesive (Surgibond, SMI AG, St. Vith, Belgium) onto slightly larger pieces of tissue gaze, which were placed into 6-or 12-well plates containing CO_2_ independent culture medium (Gibco, Thermo Fisher Scientific, Dreieich, Germany) supplemented with 0.1% [w/v] cell-culture grade BSA (Sigma Aldrich, St. Louis, Missouri) for about 10 min on ice. For inoculation, the medium was removed, a sterile piece of filter paper of 5 mm diameter was placed onto each skin sheet, and 10 µL of MNT buffer without or with varying amounts of HSV-1 were pipetted into the center of a given filter paper. After 30 min at 37°C and 5% CO_2_, the filter papers were removed, and 1 mL/well of DMEM supplemented with 10% [v/v] FBS, 100 IU/mL penicillin, 100 μg/mL streptomycin, and 1 μg/mL amphotericin B was added. The skin explants were incubated at 37°C and 5% CO_2_ for the indicated times and fixed with 3% paraformaldehyde in PBS at RT for 24 h. For quality control, images of the skin explants were documented with a wide-field fluorescence microscope (c.f. Fig. 1; ZEISS observer Z1, Carl Zeiss Microscopy GmbH Jena, Germany; CMOS camera).

For synchronous HSV-1 infections of cultured cells, Vero cells were pre-cooled for 20 min on ice and inoculated with an MOI of 10 PFU per cell or mock treated as control in CO_2_-independent medium containing 0.1% (w/v) BSA for 2 h on ice while rocking. The cells were then shifted to regular growth medium at 37°C and 5% CO_2_ for 1 h. Non-internalized virus was inactivated by a 3 min acid wash (40 mM citrate, 135 mM NaCl, 10 mM KCl, pH 3) at 4°C and after washing cells were further incubated in Vero medium for indicated times.

### Detection of host and HSV-1 genomes and transcripts

Infected skin pieces were removed from the filter papers, cut into small pieces of about 1 mm^3^, and sheared (TissueRuptor II; Qiagen, Hilden, Germany). We used the QIAamp DNA mini and Blood mini kits (Qiagen) to isolate the DNA, and the Q5 High-Fidelity DNA Polymerase (Thermo Fisher Scientific, Dreieich, Germany) with the primers HSV-1 gB-forward (5′-gtagccgtaaaacggggaca-3′) and gB-reverse (5′-ccgacctcaagtacaacccc-3′) or murine pthrp-forward (5′-ggtatctgccctcatcgtctg-3′) and pthrp-reverse (5′-cgtttcttcctccaccatctg-3′) at an annealing temperature of 60 °C to measure the amount of HSV-1 genomes relative to the host genome. We used Trizol (Thermo Fischer Scientific, Dreieich, Germany) to isolate the RNA, and LunaScript® RT SuperMix to produce the cDNA. The cDNA was amplified with Phusion^TM^ high fidelity DNA polymerase (Thermo Fisher Scientific, Dreieich, Germany) and the primers HSV-1 gB-forward (5′-cgcatcaagaccacctcctc-3′) and gB-reverse (5′-agcttgcgggcctcgtt-3′) or murine actin-forward (5′-ggctgtattcccctccatcg-3′) and actin-reverse (5′-ccagttggtaacaatgccatgt-3′) for amplification.

### One-step growth curves

Sub-confluent Vero were inoculated with a multiplicity of infection of 5 PFU per cell for 1 h at room temperature on a rocking platform and supernatants and the cells were harvested at the indicated time points. The cells were scraped in MNT buffer (30 mM MES, 100 mM NaCl, 20 mM Tris, pH 7.4) and virus was released by three cycles of freeze-thawing. Intra– and extracellular virus titers were titrated on Vero cells as described before (Grosche et al., 2019). For this, cells were inoculated with serial dilutions of the virus samples and incubated for 1 h at room temperature while rocking. Afterwards, they were incubated with growth medium containing 20 µg per ml pooled human IgGs (Sigma-Adrich, Schnelldorf, Germany). To determine the plaque size, images of living cells were acquired using a Zeiss Observer Microscope and 10x objective with appropriate filter sets and analysed in the FIJI software (version 1.50a). Plaque margins were outlined manually along the line where the CheVP26 fluorescence intensity dropped using the “freehand selection” tool. The area of the plaque was determined using the measurement function of the FIJI software. To determine the number of plaques, the cells were fixed in absolute methanol three days later and stained with 0.1% (w/v) crystal violet.

### Immunoblot

Cells were washed with PBS, lysed with hot sample buffer (50 mM Tris-HCl, pH 6.8, 1% [w/v] SDS, 1% [v/v] β-mercaptoethanol, 5% [v/v] glycerol, 0.001% [w/v] bromphenol blue) containing protease inhibitors AEL (aprotinin, E-64, leupeptin, Sigma), ABP (antipain, bestatin, pepstatin, Sigma) and PMSF (phenylmethylsulfonylfluorid in isopropanol, Roth, Karlsruhe, Germany). Cell lysates were boiled at 95°C for 5 min and homogenized by needling. The proteins were separated in by 7.5-18% SDS-PAGE and transferred in 24 mM Tris, 193 mM glycine, 0.035% [w/v] SDS and 15% [v/v] methanol to nitrocellulose membranes (Pall Corporation, Pensacola, FL, USA). After blocking with 5% [w/v] low-fat milk powder in PBS containing 0.1% [v/v] Tween 20 (PBS-T), the membranes were probed with primary antibodies and secondary antibodies coupled to fluorescent dyes (Table 2; LI-COR Biotechnology, Bad Homburg, Germany), and documented with line scanning and a digital image sensor (Odyssey Infrared Imaging System, Li-COR Biosciences, NE, USA).

### Fluorescence microscopy

Cultured cells were fixed at room temperature with 3% [w/v] paraformaldehyde for 20 min and permeabilized with 0.1% Triton-X-100 for 5 min, or treated at 37°C with 3.7% [w/v] paraformaldehyde, 0.05% [w/v] glutaraldehyde, 0.5% Triton-X-100 in PHEMO buffer with 68 mM PIPES, 25 mM HEPES, pH 6.9, 15 mM EGTA, 3 mM MgCl_2_, 10% dimethyl sulfoxide (Döhner et al., 2002). After both fixation protocols, the specimen was incubated with 50 mM NH_4_Cl for 10 min inactivate any remaining paraformaldehyde and glutaraldehyde. The HSV1 Fc-receptor and unspecific protein binding sites were blocked in blocking buffer (10% [v/v] human serum of an HSV-1 seronegative volunteer, 0.5% [w/v] BSA, in PBS, pH 7.4). The samples were labeled with primary and pre-adsorbed secondary antibodies to prevent species cross-reactivity (Table 2). The cover slips were mounted in Mowiol® 4-88 containing 2.5% (w/v) 1,4-diazabicyclo-[2.2.2]octane. Experiments were analyzed using a confocal fluorescence microscope equipped with a 63x objective (TCS SP6, LEICA Microsystems, Wetzlar, Germany).

For immunolabeling of cryosections, the ear skin explants were fixed at 4°C with 3% paraformaldehyde in PBS overnight, treated with 50 mM NH_4_Cl in PBS for 30 min to quench residual paraformaldehyde, and infiltrated with 30% [w/v] sucrose in PBS overnight at 4°C. After embedding on dry ice in O.C.T. compound (Sakura Finetek Europe B.V., Umkirch, Germany), the specimens were cut into 8 to 20 μm thick sections (Leica CM3050 S cryostat, Wetzlar, Germany) and transferred to adhesive glass slides (Thermo ScientificTM SuperFrost Ultra PlusTM GOLD, Thermo Fisher Scientific, Dreieich, Germany). The sections were incubated for 30 min at RT in blocking buffer with 0.3% [v/v] Triton-X 100, labeled with primary antibodies in blocking buffer at 4°C overnight and pre-adsorbed secondary antibodies to prevent species cross-reactivity (Table 2) in BB at RT for 90 min, and mounted in Mowiol® 4-88 containing 2.5% [w/v] 1,4-diazabicyclo-[2.2.2]octane. Images of single optical sections of about 0.5 μm thickness were captured with a confocal microscope and a 20x or 63x objective (Leica inverted 3).

Images of a given experiment were adjusted with identical linear contrast and brightness settings, merged, and evaluated using the Fiji image analysis platform (Schindelin et al., 2012)

### 2-Photon fluorescence microscopy

Ear sheets were fixed in 3% paraformaldehyde in PBS at 4°C overnight and attached to the bottom of an imaging chamber with tissue adhesive. The samples were investigated at RT with 2-Photon imaging using two laser lines in parallel and PBS as the optical medium (LaVision Bio Tec TrimpScope 2; Miltenyi Biotec, Bielefeld, Germany). mCherry was excited with a Spectra-Physics Insight laser set to 1105 nm and GFP and dermal collagen were visualized by a second laser set to 910 nm at an intensity of 1 to 10% of its maximum power. The emitted light was captured with a 20x objective with a standard voxel size of 4 μm^3^ (NA 0.9; ZEISS) and recorded with photononic multiplier tubes (Hamamatsu) with different bandpass filters; namely a blue channel at 400-450 nm for the dermal collagen, a green channel at 510-550 nm for GFP, and a red channel at 610-640 nm for mCherry. Images were acquired with the LaVision Inspector Software Version 3 and analyzed with the Imaris software (Version 9.1; Bitplane at Oxford Instruments; Oxfordshire, UK) as reported before (Halle et al., 2016).

### Histopathology and immunohistochemistry

Skin explants were fixed in 3% paraformaldehyde in PBS overnight at 4°C, embedded in paraffin after trimming of cross sections and cut into 5 μm thick sections. For histopathological examination, sections were stained with hematoxylin and eosin. The following criteria have been determined for grading the severity of epithelial hyperplasia: mild (up to 5 layers of epithelial cells), moderate (6-10 layers of epithelial cells), marked (more than 10 layers of epithelial cells). Immunohistochemistry was performed as described previously (Störk et al., 2021). Briefly, the sections were deparaffinized and rehydrated followed by inhibition of endogenous peroxidase activity. Afterwards, antigen retrieval was achieved by incubation of sections with citrate buffer (pH 6.0) in the microwave at 800 W for 20 min. Primary antibodies were applied in their respective concentration (pan-cytokeratin AE1/AE3: 1:500, HSV-1 ICP8: 1:1600, diluted in PBS with addition of 1% bovine serum albumin) overnight at 4°C. Negative controls were generated by the substitution of primary antibodies with Balb/c mouse ascites fluid (diluted 1:1000 in PBS). Thereafter, the sections were incubated with EnVision+ System-HRP labeled polymer (Dako Agilent Pathology Solutions) for 30 min at room temperature. Visualization of immunpositive reactions was achieved by the application of 3,3’-diaminobenzidine tetrahydrochloride (0.05%, Sigma Aldrich Chemie GmbH, Darmstadt, Germany) with 0.03% hydrogen peroxide for 5 min at room temperature. Finally, sections were counterstained with Mayeŕs hematoxylin (Roth C. GmbH & Co KG, Karlsruhe, Germany).

### Electron microscopy

Ear sheets were infected with HSV1-Che at 1 x 10^7^ PFU per filter paper for 48 h. The samples were fixed in 2% [w/v] glutaraldehyde, 2.5% [w/v] paraformaldehyde, 165 mM sodium cacodylate, pH 7.4, 2 mM CaCl_2_, and 10 mM MgCl_2_ at RT for 1 h. To locate the small infection centers, the explants were placed with the apical surface onto coverslips with a gridded checkerboard of alphanumeric patterns (P35G-2-14-C; MatTek Europe, Slovak Republic), and embedded in 10% [w/v] gelatin in PBS at 37°C. The coverslips were immediately placed on an ice-cold metal plate to solidify the gelatin, and fresh fixative was added on ice for 1 h to crosslink the gelatin to the skin sheets. As ethanol and propylenoxide used for resin embedding destroy the fluorescence of mCherry, we localized infection centers at this point of the protocol in reference to the gridded checkerboard by their HSV-1 mediated mChe expression using a wide-field fluorescence microscope (Leica Microsystems, Germany), and the filter set N2.1 containing the emission long pass filter LP590 which largely blocks the autofluorescence generated by the glutaraldehyde fixation. Subsequently, the specimen were contrasted with 1% [w/v] OsO_4_ in 165 mM cacodylate buffer, pH 7.4 containing 1.5% [w/v] K_3_[Fe(CN)_6_] for 1 h, followed by 0.5% [w/v] uranyl acetate in 50% (v/v) ethanol overnight, the ethanol chain for dehydration, a propylenoxide intermediate, and embedding in expoxy resin (Epon, Serva, Heidelberg, Germany). The gridded coverslips were detached from the polymerized resin but had left their checkerboard imprints in the resin to locate the positions of the infection centers. These skin areas were carved from the resin, cut in half, and each half was glued onto a separate resin block at a 90° angle with the cut surface facing the microtome knife to obtain cross-sections through the infected skin explants. Ultrathin sections of 50-nm thickness were cut and further contrasted using lead citrate (Reynolds, 1963) and uranyl acetate (Watson, 1958). Images were taken at 80 kV with a transmission electron microscope (Morgani, FEI, The Netherlands) equipped with a digital camera (Veleta; Olympus Soft Imaging Solutions, Germany) and processed using the Fiji image analysis platform (Schindelin et al., 2012)

### Statistical analyses

Data were presented in scatter plots with lines at the median with the median interquartile range as error bars. We performed Wilcoxon signed-rank, Mann-Whitney U, and t-tests (GraphPad Prism v5.0; La Jolla, California), and calculated the standardized effect sizes and the statistical power (Faul et al., 2007; Gpower 3.1; https://gpower.software.informer.com/3.1/). A p-value of less than 0.05 was considered significant.

### Ethics

The mouse experiments were approved by the local Advisory Committee on Institutional Animal Care and Research (MHH) and the Lower Saxony State Office for Consumer Protection and Food Safety (LAVES; *Landesamt für and Verbraucherschutz und Lebensmittelsicherheit,* Hannover, Germany) through license §4 2018/189 of the German Animal Welfare Law (*Tierschutzgesetz*), and conducted according to the German animal welfare law, the German regulations of the Society for Laboratory Animal Science (GV-SOLAS), the European Directive 2010/63/EU, and the European Health Law of the Federation of Laboratory Animal Science Association (FELASA).

### Author contributions

TR, BS, and SH wrote the manuscript and conceived the study. TR, XL, KD, ABu, ABi, AP, MdlR, RB, BS, and SH performed experiments, data curation, and formal analysis. TR, WB, RF, BS and SH acquired grant funding. All authors reviewed and edited the manuscript prior to submission.

## Acknowledgements

We are grateful to Abel Viejo Borbolla, Julio Villalvazo Guerrero, and Manutea Serrero for many constructive discussions and feedback, Michaela Cappucci for support in characterizing the HSV1-Che26-VP11/12GFP strains (Institute of Virology, Hannover Medical School). We thank Julia Baskas, Petra Grünig, Jana-Svea Harre, Claudia Herrmann, Kerstin Rohn, Kerstin Schöne, Caroline Schütz and Danuta Waschke for their technical support (Department of Pathology, University of Veterniary Medicine Hannover). For providing the clinical isolate HSV-1 strain 17^+^, we are grateful to the late John Subak-Sharpe (MRC, Glasgow, United Kingdom), and for donating essential antibodies, we thank Helena Browne and Tony Minson (anti-VP16; University of Cambridge, UK), Gillian Elliott (anti-VP22; University of Surrey, UK), and Michael Nassal (anti-GFP; University Hospital Freiburg, Germany).

Our work was funded by the *Deutsche Forschungsgemeinschaft* (DFG EXC2155 RESIST, grant ID 390874280 to RF, TW and BS; DFG CRC 900, grant ID 158989968, project B1 to RF, project B3 to SH, and project C2 to BS; FOR 2830 to RF, FO 334/7-2), the Niedersachsen-Research Network on Neuroinfectiology of the Ministry of Science and Culture (N-RENNT to BS, RF, and WB), and the German Center for Infection Research (DZIF, TTU Infections of the immunocompromised host to BS). TR was a PhD fellow from the German Academic Exchange Service (DAAD, https://www.daad.de/en/; grant ID 91727753), MdlR was supported by the DFG RTN 2485 VIPER (ID grant 398066876). TR and AB were supported by the Hannover Biomedical Research School (HBRS) and its Center for Infection Biology (ZIB). We are grateful to the Research Core Unit for Laser Microscopy (ReCoLa) and the Research Core Unit for Electron Microscopy at Hannover Medical School for their support.

## Abbreviations

BAC: bacterial artificial chromosome
BSA: bovine serum albumin
dpi: day post infection
gB: glycoprotein B
gD: glycoprotein D
hpi: hour post infection
HSV-1: herpes simplex virus type 1
ICP: infected cell protein
MOI: multiplicity of infection
PFU: plaque forming unit
VP: viral protein

## REFERENCES

1. Aho V, Salminen S, Mattola S, Gupta A, Flomm F, Sodeik B, Bosse JB, Vihinen-Ranta M. 2021. Infection-induced chromatin modifications facilitate translocation of herpes simplex virus capsids to the inner nuclear membrane. PLoS Pathog 17:e1010132. doi:10.1371/journal.ppat.1010132

2. Allan RS, Smith CM, Belz GT, van Lint AL, Wakim LM, Heath WR, Carbone FR. 2003. Epidermal viral immunity induced by CD8 dendritic cells but not by Langerhans cells. Science 301:1925–8. doi:10.1126/science.1087576

3. Amberg N, Holcmann M, Stulnig G, Glitzner E, Sibilia M. 2017. Effects of depilation methods on imiquimod-induced skin inflammation in mice. Journal of Investigative Dermatology 137:528–531. doi:10.1016/j.jid.2016.09.018

4. Bedoui S, Whitney PG, Waithman J, Eidsmo L, Wakim L, Caminschi I, Allan RS, Wojtasiak M, Shortman K, Carbone FR, Brooks AG, Heath WR. 2009. Cross-presentation of viral and self antigens by skin-derived CD103+ dendritic cells. Nat Immunol 10:488–95. doi:10.1038/ni.1724

5. Bertram KM, Truong NR, Smith JB, Kim M, Sandgren KJ, Feng KL, Herbert JJ, Rana H, Danastas K, Miranda-Saksena M, Rhodes JW, Patrick E, Cohen RC, Lim J, Merten SL, Harman AN, Cunningham AL. 2021. Herpes Simplex Virus type 1 infects langerhans cells and the novel epidermal dendritic cell, Epi-cDC2s, via different entry pathways. PLoS Pathog 17:e1009536. doi:10.1371/journal.ppat.1009536

6. Bodda C, Reinert LS, Fruhwürth S, Richardo T, Sun C, Zhang B-C, Kalamvoki M, Pohlmann A, Mogensen TH, Bergström P, Agholme L, O’Hare P, Sodeik B, Gyrd-Hansen M, Zetterberg H, Paludan SR. 2020. HSV1 VP1-2 deubiquitinates STING to block type I interferon expression and promote brain infection. J Exp Med 217. doi:10.1084/jem.20191422

7. Bosse JB, Hogue IB, Feric M, Thiberge SY, Sodeik B, Brangwynne CP, Enquist LW. 2015. Remodeling nuclear architecture allows efficient transport of herpesvirus capsids by diffusion. Proceedings of the National Academy of Sciences 112. doi:10.1073/pnas.1513876112

8. Brown SM, Ritchie DA, Subak-Sharpe JH. 1973. Genetic Studies with Herpes Simplex Virus Type 1. The Isolation of temperature-sensitive mutants, their arrangement into complementation groups and recombination analysis leading to a linkage map. Journal of General Virology 18:329–346. doi:10.1099/0022-1317-18-3-329

9. Buch A, Müller O, Ivanova L, Döhner K, Bialy D, Bosse JB, Pohlmann A, Binz A, Hegemann M, Nagel C-H, Koltzenburg M, Viejo-Borbolla A, Rosenhahn B, Bauerfeind R, Sodeik B. 2017. Inner tegument proteins of Herpes Simplex Virus are sufficient for intracellular capsid motility in neurons but not for axonal targeting. PLoS Pathog 13:e1006813. doi:10.1371/journal.ppat.1006813

10. Chopra S, Roesner LM, Döhner K, Zeitvogel J, Traidl S, Rodriguez E, Harder I, Wolfgang L, Weidinger S, Schulz TF, Sodeik B, Werfel T. 2024. Identification and characterization of collagen XXIII alpha I as a novel risk factor for eczema herpeticum. (Preprint-MedRvix)

11. Coates M, Blanchard S, MacLeod AS. 2018. Innate antimicrobial immunity in the skin: A protective barrier against bacteria, viruses, and fungi. PLoS Pathog 14:e1007353. doi:10.1371/journal.ppat.1007353

12. Cohen JI. 2020. Herpesvirus latency. J Clin Invest 130:3361–3369. doi:10.1172/JCI136225

13. Cunningham AL, Turner RR, Miller AC, Para MF, Merigan TC. 1985. Evolution of recurrent herpes simplex lesions. An immunohistologic study. J Clin Invest 75:226–33. doi:10.1172/JCI111678

14. De Bernardo R. 1992. Herpes gladiatorum. N Engl J Med 326:647–8. doi:10.1056/nejm199202273260919

15. De La Cruz NC, Möckel M, Wirtz L, Sunaoglu K, Malter W, Zinser M, Knebel-Mörsdorf D. 2021. Ex Vivo infection of human skin with herpes simplex virus 1 reveals mechanical wounds as insufficient entry portals via the skin surface. J Virol 95:e0133821. doi:10.1128/JVI.01338-21

16. Döhner K, Radtke K, Schmidt S, Sodeik B. 2006. Eclipse Phase of Herpes Simplex Virus Type 1 Infection: Efficient dynein-mediated capsid transport without the small capsid protein VP26. J Virol 80:8211–8224. doi:10.1128/JVI.02528-05

17. Döhner K, Ramos-Nascimento A, Bialy D, Anderson F, Hickford-Martinez A, Rother F, Koithan T, Rudolph K, Buch A, Prank U, Binz A, Hugel S, Lebbink RJRJ, Hoeben RCRC, Hartmann E, Bader M, Bauerfeind R, Sodeik B, Hu S. 2018. Importin α1 is required for nuclear import of herpes simplex virus proteins and capsid assembly in fibroblasts and neurons. PLoS Pathog 14:e1006823. doi:10.1371/journal.ppat.1006823

18. Döhner K, Serrero MC, Viejo-Borbolla A, Sodeik B. 2024. A hitchhiker’s guide through the cell: the world according to the capsids of alphaherpesviruses. Annu Rev Virol. doi:10.1146/annurev-virology-100422-022751

19. Döhner K, Wolfstein A, Prank U, Echeverri C, Dujardin D, Vallee R, Sodeik B. 2002. Function of dynein and dynactin in herpes simplex virus capsid transport. Mol Biol Cell 13:2795–2809. doi:10.1091/mbc.01-07-0348

20. Faul F, Erdfelder E, Lang A-G, Buchner A. 2007. G*Power 3: A flexible statistical power analysis program for the social, behavioral, and biomedical sciences. Behav Res Methods 39:175–191. doi:10.3758/BF03193146

21. Feder HM. 1983. Herpetic Whitlow. American Journal of Diseases of Children 137:861. doi:10.1001/archpedi.1983.02140350035009

22. Gebhardt T, Whitney PG, Zaid A, Mackay LK, Brooks AG, Heath WR, Carbone FR, Mueller SN. 2011. Different patterns of peripheral migration by memory CD4+ and CD8+ T cells. Nature 477:216–219. doi:10.1038/nature10339

23. Gnann JW, Whitley RJ. 2017. Herpes Simplex Encephalitis: an Update. Curr Infect Dis Rep 19:13. doi:10.1007/s11908-017-0568-7

24. Grosche L, Döhner K, Düthorn A, Hickford-Martinez A, Steinkasserer A, Sodeik B. 2019. Herpes Simplex Virus Type 1 propagation, titration and single-step growth curves. Bio Protoc 9. doi:10.21769/BioProtoc.3441

25. Halle S, Keyser KA, Stahl FR, Busche A, Marquardt A, Zheng X, Galla M, Heissmeyer V, Heller K, Boelter J, Wagner K, Bischoff Y, Martens R, Braun A, Werth K, Uvarovskii A, Kempf H, Meyer-Hermann M, Arens R, Kremer M, Sutter G, Messerle M, Förster R. 2016. In vivo killing capacity of cytotoxic T cells is limited and involves dynamic interactions and T cell cooperativity. Immunity 44:233–45. doi:10.1016/j.immuni.2016.01.010

26. Harbour DA, Hill TJ, Blyth WA. 1981. Acute and recurrent herpes simplex in several strains of mice. Journal of General Virology 55:31–40. doi:10.1099/0022-1317-55-1-31

27. Hill TJ, Field HJ, Blyth WA. 1975. Acute and recurrent Infection with herpes simplex virus in the mouse: a model for studying latency and recurrent disease. Journal of General Virology 28:341–353. doi:10.1099/0022-1317-28-3-341

28. Hill TJ, Shimeld C. 1998. Models of recurrent infection with HSV in the skin and eye of the mouse herpes simplex virus protocols. New Jersey: Humana Press. pp. 273–290. doi:10.1385/0-89603-347-3:273

29. Hor JL, Whitney PG, Zaid A, Brooks AG, Heath WR, Mueller SN. 2015. Spatiotemporally distinct interactions with dendritic cell subsets facilitates CD4+ and CD8+ T Cell activation to localized viral infection. Immunity 43:554–565. doi:10.1016/j.immuni.2015.07.020

30. Hussain MT, Stanfield BA, Bernstein DI. 2024. Small animal models to study herpes simplex virus infections. Viruses 16:1037. doi:10.3390/v16071037

31. Ivanova L, Buch A, Döhner K, Pohlmann A, Binz A, Prank U, Sandbaumhüter M, Bauerfeind R, Sodeik B. 2016. Conserved tryptophan motifs in the large tegument protein pUL36 are required for efficient secondary envelopment of herpes simplex virus capsids. J Virol 90:5368–5383. doi:10.1128/JVI.03167-15

32. James C, Harfouche M, Welton NJ, Turner KM, Abu-Raddad LJ, Gottlieb SL, Looker KJ. 2020. Herpes simplex virus: global infection prevalence and incidence estimates, 2016. Bull World Health Organ 98:315–329. doi:10.2471/BLT.19.237149

33. Jones J, Depledge DP, Breuer J, Ebert-Keel K, Elliott G. 2019. Genetic and phenotypic intrastrain variation in herpes simplex virus type 1 Glasgow strain 17 syn+-derived viruses. Journal of General Virology 100:1701–1713. doi:10.1099/jgv.0.001343

34. Kim M, Osborne NR, Zeng W, Donaghy H, McKinnon K, Jackson DC, Cunningham AL. 2012. Herpes simplex virus antigens directly activate NK Cells via TLR2, thus facilitating their presentation to CD4 T Lymphocytes. The Journal of Immunology 188:4158– 4170. doi:10.4049/jimmunol.1103450

35. Kim M, Truong NR, James V, Bosnjak L, Sandgren KJ, Harman AN, Nasr N, Bertram KM, Olbourne N, Sawleshwarkar S, McKinnon K, Cohen RC, Cunningham AL. 2015. Relay of herpes simplex virus between Langerhans cells and dermal dendritic Cells in human skin. PLoS Pathog 11:e1004812. doi:10.1371/journal.ppat.1004812

36. Kollias CM, Huneke RB, Wigdahl B, Jennings SR. 2015. Animal models of herpes simplex virus immunity and pathogenesis. J Neurovirol 21:8–23. doi:10.1007/s13365-014-0302-2

37. Kopfnagel V, Dreyer S, Baumert K, Stark M, Harder J, Hofmann K, Kleine M, Buch A, Sodeik B, Werfel T. 2020. RNase 7 promotes sensing of self-DNA by human keratinocytes and activates an antiviral immune response. Journal of Investigative Dermatology 140:1589–1598.e3. doi:10.1016/j.jid.2019.09.029

38. Koyuncu OO, Enquist LW, Engel EA. 2021. Invasion of the nervous system. Curr Issues Mol Biol 41:1–62. doi:10.21775/cimb.041.001

39. Krawczyk A, Dirks M, Kasper M, Buch A, Dittmer U, Giebel B, Wildschütz L, Busch M, Goergens A, Schneweis KE, Eis-Hübinger AM, Sodeik B, Heiligenhaus A, Roggendorf M, Bauer D. 2015. Prevention of herpes simplex virus induced stromal keratitis by a glycoprotein B-specific monoclonal antibody. PLoS One 10:e0116800. doi:10.1371/journal.pone.0116800

40. Lin, A. E., Greco, T. M., Döhner, K., Sodeik, B., & Cristea, I. M. (2013). A proteomic perspective of inbuilt viral protein regulation: pUL46 tegument protein is targeted for degradation by ICP0 during herpes simplex virus type 1 infection. Molecular & Cellular Proteomics, 12(11), 3237–3252. 10.1074/mcp.M113.030866

41. Lee J-N, Jee S-H, Chan C-C, Lo W, Dong C-Y, Lin S-J. 2008. The effects of depilatory agents as penetration enhancers on human stratum corneum structures. J Invest Dermatol 128:2240–7. doi:10.1038/jid.2008.82

42. Looker KJ, Magaret AS, Turner KME, Vickerman P, Gottlieb SL, Newman LM. 2015. Global estimates of prevalent and incident herpes simplex virus type 2 infections in 2012. PLoS One 10:e114989. doi:10.1371/journal.pone.0114989

43. Mackay LK, Stock AT, Ma JZ, Jones CM, Kent SJ, Mueller SN, Heath WR, Carbone FR, Gebhardt T. 2012. Long-lived epithelial immunity by tissue-resident memory T (TRM) cells in the absence of persisting local antigen presentation. Proceedings of the National Academy of Sciences 109:7037–7042. doi:10.1073/pnas.1202288109

44. Menon GK, Cleary GW, Lane ME. 2012. The structure and function of the stratum corneum. Int J Pharm 435:3–9. doi:10.1016/j.ijpharm.2012.06.005

45. Möckel M, De La Cruz NC, Rübsam M, Wirtz L, Tantcheva-Poor I, Malter W, Zinser M, Bieber T, Knebel-Mörsdorf D. 2022. Herpes Simplex Virus 1 can bypass impaired epidermal barriers upon *Ex Vivo* infection of skin from atopic dermatitis patients. J Virol 96. doi:10.1128/jvi.00864-22

46. Nagel C-H, Dohner K, Fathollahy M, Strive T, Borst EM, Messerle M, Sodeik B. 2008. Nuclear egress and envelopment of herpes simplex virus capsids analyzed with dual-color fluorescence HSV1(17+). J Virol 82:3109–3124. doi:10.1128/JVI.02124-07

47. Nagel C-H, Pohlmann A, Sodeik B. 2014. Construction and characterization of bacterial artificial chromosomes (BACs) containing herpes simplex virus full-length genomes. Methods Mol Biol 1144:43–62. doi:10.1007/978-1-4939-0428-0_4

48. Nestle FO, Di Meglio P, Qin J-Z, Nickoloff BJ. 2009. Skin immune sentinels in health and disease. Nat Rev Immunol 9:679–691. doi:10.1038/nri2622

49. Ojala PM, Sodeik B, Ebersold MW, Kutay U, Helenius A. 2000. Herpes simplex virus type 1 entry into host cells: reconstitution of capsid binding and uncoating at the nuclear pore complex in vitro. Mol Cell Biol 20:4922–31. doi:10.1128/MCB.20.13.4922-4931.2000

50. Oyoshi MK, Larson RP, Ziegler SF, Geha RS. 2010. Mechanical injury polarizes skin dendritic cells to elicit a T(H)2 response by inducing cutaneous thymic stromal lymphopoietin expression. J Allergy Clin Immunol 126:976–84, 984.e1–5. doi:10.1016/j.jaci.2010.08.041

51. Pasparakis M, Haase I, Nestle FO. 2014. Mechanisms regulating skin immunity and inflammation. Nat Rev Immunol 14:289–301. doi:10.1038/nri3646

52. Pellett PE, Kousoulas KG, Pereira L, Roizman B. 1985. Anatomy of the herpes simplex virus 1 strain F glycoprotein B gene: primary sequence and predicted protein structure of the wild type and of monoclonal antibody-resistant mutants. J Virol 53:243–53. doi:10.1128/JVI.53.1.243-253.1985

53. Petermann P, Thier K, Rahn E, Rixon FJ, Bloch W, Özcelik S, Krummenacher C, Barron MJ, Dixon MJ, Scheu S, Pfeffer K, Knebel-Mörsdorf D. 2015. Entry mechanisms of herpes simplex virus 1 into murine epidermis: involvement of Nectin-1 and herpesvirus entry mediator as cellular receptors. J Virol 89:262–274. doi:10.1128/JVI.02917-14

54. Piret J, Boivin G. 2020. Immunomodulatory strategies in herpes simplex virus encephalitis. Clin Microbiol Rev 33. doi:10.1128/CMR.00105-19

55. Proença JT, Nelson D, Nicoll MP, Connor V, Efstathiou S. 2016. Analyses of herpes simplex virus type 1 latency and reactivation at the single cell level using fluorescent reporter mice. Journal of General Virology 97:767–777. doi:10.1099/jgv.0.000380

56. Puttur FK, Fernandez MA, White R, Roediger B, Cunningham AL, Weninger W, Jones CA. 2010. Herpes Simplex Virus Infects Skin γδ T Cells before Langerhans Cells and impedes migration of infected Langerhans cells by inducing apoptosis and blocking E-Cadherin downregulation. The Journal of Immunology 185:477–487. doi:10.4049/jimmunol.0904106

57. Rahn E, Petermann P, Thier K, Bloch W, Morgner J, Wickström SA, Knebel-Mörsdorf D. 2015. Invasion of herpes simplex virus type 1 into murine epidermis: an ex vivo infection study. Journal of Investigative Dermatology 135:3009–3016. doi:10.1038/jid.2015.290

58. Rahn E, Thier K, Petermann P, Rübsam M, Staeheli P, Iden S, Niessen CM, Knebel-Mörsdorf D. 2017. Epithelial barriers in murine skin during herpes simplex virus 1 infection: the role of tight junction formation. Journal of Investigative Dermatology 137:884–893. doi:10.1016/j.jid.2016.11.027

59. Reichert MN, Koewler NJ, Hargis AM, Felgenhauer JL, Impelluso LC. 2023. Effects of depilatory cream formulation and contact time on mouse skin. Journal of the American Association for Laboratory Animal Science 62:153–162. doi:10.30802/AALAS-JAALAS-22-000065

60. Reynolds ES. 1963. The use of lead citrate at high pH as an electron-opaque stain in electron microscopy. J Cell Biol 17:208–212. doi:10.1083/jcb.17.1.208

61. Sandbaumhüter M, Döhner K, Schipke J, Binz A, Pohlmann A, Sodeik B, Bauerfeind R. 2013. Cytosolic herpes simplex virus capsids not only require binding inner tegument protein pUL36 but also pUL37 for active transport prior to secondary envelopment. Cell Microbiol 15:248–69. doi:10.1111/cmi.12075

62. Sawtell NM, Thompson RL. 2014. Herpes simplex virus mutant generation and dual-detection methods for gaining insight into latent/lytic cycles in vivo. pp. 129–147. doi:10.1007/978-1-4939-0428-0_9

63. Schindelin J, Arganda-Carreras I, Frise E, Kaynig V, Longair M, Pietzsch T, Preibisch S, Rueden C, Saalfeld S, Schmid B, Tinevez J-Y, White DJ, Hartenstein V, Eliceiri K, Tomancak P, Cardona A. 2012. Fiji: an open-source platform for biological-image analysis. Nat Methods 9:676–82. doi:10.1038/nmeth.2019

64. Serup J. 2023. Technical and clinical complications of cosmetic tattooing. pp. 225–244. doi:10.1159/000526048

65. Silva JR, Lopes AH, Talbot J, Cecilio NT, Rossato MF, Silva RL, Souza GR, Silva CR, Lucas G, Fonseca BA, Arruda E, Alves-Filho JC, Cunha FQ, Cunha TM. 2017. Neuroimmune–Glia interactions in the sensory ganglia account for the development of acute herpetic neuralgia. The Journal of Neuroscience 37:6408–6422. doi:10.1523/JNEUROSCI.2233-16.2017

66. Singh M, Goodyear HM, Breuer J. 2019. Herpes Simplex Virus infections Harper’s Textbook of Pediatric Dermatology. Wiley. pp. 598–611. doi:10.1002/9781119142812.ch50

67. Snijder B, Sacher R, Rämö P, Liberali P, Mench K, Wolfrum N, Burleigh L, Scott CC, Verheije MH, Mercer J, Moese S, Heger T, Theusner K, Jurgeit A, Lamparter D, Balistreri G, Schelhaas M, De Haan CAM, Marjomäki V, Hyypiä T, Rottier PJM, Sodeik B, Marsh M, Gruenberg J, Amara A, Greber U, Helenius A, Pelkmans L. 2012. Single-cell analysis of population context advances RNAi screening at multiple levels. Mol Syst Biol 8:579. doi:10.1038/msb.2012.9

68. Sodeik B, Ebersold MW, Helenius A. 1997. Microtubule-mediated transport of incoming herpes simplex virus 1 capsids to the nucleus. Journal of Cell Biology 136:1007–1021. doi:10.1083/jcb.136.5.1007

69. St Laurent G, Toma I, Seilheimer B, Cesnulevicius K, Schultz M, Tackett M, Zhou J, Ri M, Shtokalo D, Antonets D, Jepson T, McCaffrey TA. 2021. RNAseq analysis of treatment-dependent signaling changes during inflammation in a mouse cutaneous wound healing model. BMC Genomics 22:854. doi:10.1186/s12864-021-08083-2

70. Stock AT, Mueller SN, van Lint AL, Heath WR, Carbone FR. 2004. Cutting edge: prolonged antigen presentation after herpes simplex virus-1 skin infection. The Journal of Immunology 173:2241–2244. doi:10.4049/jimmunol.173.4.2241

71. Störk, T., de le Roi, M., Haverkamp, A.-K., Jesse, S. T., Peters, M., Fast, C., Gregor, K. M., Könenkamp, L., Steffen, I., Ludlow, M., Beineke, A., Hansmann, F., Wohlsein, P., Osterhaus, A. D. M. E., & Baumgärtner, W. (2021). Analysis of avian Usutu virus infections in Germany from 2011 to 2018 with focus on dsRNA detection to demonstrate viral infections. Scientific Reports, 11(1), 24191. 10.1038/s41598-021-03638-5

72. Tajpara P, Mildner M, Schmidt R, Vierhapper M, Matiasek J, Popow-Kraupp T, Schuster C, Elbe-Bürger A. 2019. A preclinical model for studying herpes simplex virus infection. J Invest Dermatol 139:673–682. doi:10.1016/j.jid.2018.08.034

73. Tan XYD, Wiseman T, Betihavas V. 2022. Risk factors for nosocomial infections and/or sepsis in adult burns patients: An integrative review. Intensive Crit Care Nurs 73:103292. doi:10.1016/j.iccn.2022.103292

74. Thier K, Petermann P, Rahn E, Rothamel D, Bloch W, Knebel-Mörsdorf D. 2017. Mechanical barriers restrict invasion of herpes simplex virus 1 into human oral mucosa. J Virol 91:JVI.01295–17. doi:10.1128/JVI.01295-17

75. Tischer BK, von Einem J, Kaufer B, Osterrieder N. 2006. Two-step red-mediated recombination for versatile high-efficiency markerless DNA manipulation in Escherichia coli. Biotechniques 40:191–7. doi:10.2144/000112096

76. Traidl S, Heinrich L, Siegels D, Rösner L, Haufe E, Harder I, Abraham S, Ertner K, Kleinheinz A, Schäkel K, Wollenberg A, Effendy I, Quist S, Asmussen A, Wildberger J, Weisshaar E, Wiemers F, Brücher J, Weidinger S, Schmitt J, Werfel T. 2023. High recurrence rate of eczema herpeticum in moderate/severe atopic dermatitis –TREAT Germany registry analysis. JDDG: Journal der Deutschen Dermatologischen Gesellschaft 21:1490–1498. doi:10.1111/ddg.15205

77. Traidl S, Roesner L, Zeitvogel J, Werfel T. 2021. Eczema herpeticum in atopic dermatitis. Allergy 76:3017–3027. doi:10.1111/all.14853

78. Tsalenchuck Y, Tzur T, Steiner I, Panet A. 2014. Different modes of herpes simplex virus type 1 spread in brain and skin tissues. J Neurovirol 20:18–27. doi:10.1007/s13365-013-0224-4

79. van Lint A, Ayers M, Brooks AG, Coles RM, Heath WR, Carbone FR. 2004. Herpes simplex virus-specific CD8+ T cells can clear established lytic infections from skin and nerves and can partially limit the early spread of virus after cutaneous inoculation. The Journal of Immunology 172:392–397. doi:10.4049/jimmunol.172.1.392

80. van Lint AL, Kleinert L, Clarke SRM, Stock A, Heath WR, Carbone FR. 2005. Latent infection with herpes simplex virus is associated with ongoing CD8+ Tcell stimulation by parenchymal cells within sensory ganglia. J Virol 79:14843–14851. doi:10.1128/JVI.79.23.14843-14851.2005

81. Wakim LM, Waithman J, van Rooijen N, Heath WR, Carbone FR. 2008. Dendritic cell-induced memory T cell activation in nonlymphoid tissues. Science 319:198–202. doi:10.1126/science.1151869

82. Wallace L, Roberts-Thompson L, Reichelt J. 2012. Deletion of K1/K10 does not impair epidermal stratification but affects desmosomal structure and nuclear integrity. J Cell Sci. doi:10.1242/jcs.097139

83. Watson ML. 1958. Staining of tissue sections for electron microscopy with heavy metals. J Cell Biol 4:475–478. doi:10.1083/jcb.4.4.475

84. Weidinger S, Beck LA, Bieber T, Kabashima K, Irvine AD. 2018. Atopic dermatitis. Nat Rev Dis Primers 4:1. doi:10.1038/s41572-018-0001-z

85. Weller SK, Lee KJ, Sabourin DJ, Schaffer PA. 1983. Genetic analysis of temperature-sensitive mutants which define the gene for the major herpes simplex virus type 1 DNA-binding protein. J Virol 45:354–66. doi:10.1128/JVI.45.1.354-366.1983

86. Wen S, Ye L, Wang X, Liu D, Yang B, Man M-Q. 2022. Aged and young mice differentially respond to tape-stripping in epidermal gene expression. Exp Dermatol 31:312–319. doi:10.1111/exd.14463

87. Whitley RJ, Roizman B. 2016. Herpes Simplex Viruses Clinical Virology. Washington, DC, USA: ASM Press. pp. 415–445. doi:10.1128/9781555819439.ch20

88. Willard M. 2002. Rapid directional translocations in virus replication. J Virol 76:5220–5232. doi:10.1128/JVI.76.10.5220-5232.2002

89. Wurzer P, Cole MR, Clayton RP, Hundeshagen G, Nunez Lopez O, Cambiaso-Daniel J, Winter R, Branski LK, Hawkins HK, Finnerty CC, Herndon DN, Lee JO. 2017. Herpesviradae infections in severely burned children. Burns 43:987–992. doi:10.1016/j.burns.2017.01.032

90. Zeitvogel J, Döhner K, Klug I, Richardo T, Sodeik B, Werfel T. 2024. Short-form thymic stromal lymphopoietin (sfTSLP) restricts herpes simplex virus infection of human primary keratinocytes. J Med Virol 96. doi:10.1002/jmv.29865

